# Discovery of a novel antifungal compound, ilicicolin K, through genetic activation of the ilicicolin biosynthetic pathway in *Trichoderma reesei*

**DOI:** 10.1101/2024.06.20.599701

**Authors:** Isabella Burger, Matthias Schmal, Kathrin Peikert, Lukas Fourtis, Christoph Suster, Christian Stanetty, Dominik Schnalzer, Ruth Birner-Gruenberger, Robert L Mach, Astrid R. Mach-Aigner, Matthias Schittmayer, Christian Zimmermann

## Abstract

In the quest to discover novel antifungal agents and new antifungal production processes, we investigated the biosynthetic gene cluster (BGC) for ilicicolin H in the fungus *Trichoderma reesei*. While the BGC is silent under standard cultivation conditions, we achieved to activate it by over-expressing its transcription factor TriliR. Successful BGC activation was confirmed by RT-qPCR, proteomic and metabolomic analyses. Metabolomic profiling upon BGC expression revealed high-yield production of the supposed main product ilicicolin H. To elucidate the functionality of this BGC, we employed a combination of overexpression and deletions of individual biosynthetic gene cluster constituents. Deletion of *triliA*, encoding for the core polyketide synthase TriliA, completely ceased product formation, as expected. In contrast to previous heterologous expression experiments, we could demonstrate that the epimerase TriliE is necessary for the formation of ilicicolin H in the native host. While we hardly observed any of the previously reported side- or shunt products associated with heterologous ilicicolin H expression, we discovered a novel member of the ilicicolin family using a metabolomic molecular networking approach. This new compound, which we termed ilicicolin K, is expressed in substantial amounts in the genetically engineered *Trichoderma reesei*, enabling us to elucidate its structure by NMR. The structure of ilicicolin K is similar to that of ilicicolin H but differs by an additional hydroxylation and an intramolecular etherification of the hydroxyl group at the pyridone towards the tyrosine moiety of the molecule. Initial tests of ilicicolin K showed antifungal activity against *Saccharomyces cerevisiae* and *Aspergillus nidulans* with a similar minimum inhibitory concentration as ilicicolin H.

## 1. Introduction

Since ancient times, mankind has valued natural products for their therapeutic potential. While plants have long been recognized for their medicinal value, fungi, although often overlooked, harbor a remarkably diverse reservoir of potentially beneficial compounds. Among them is the natural product ilicicolin H (Figure 1C **(1)**), which was discovered in 1971 as an antibiotic from the imperfect fungus *Cylindrocladium ilicicola*^1^.

**Figure 1.**
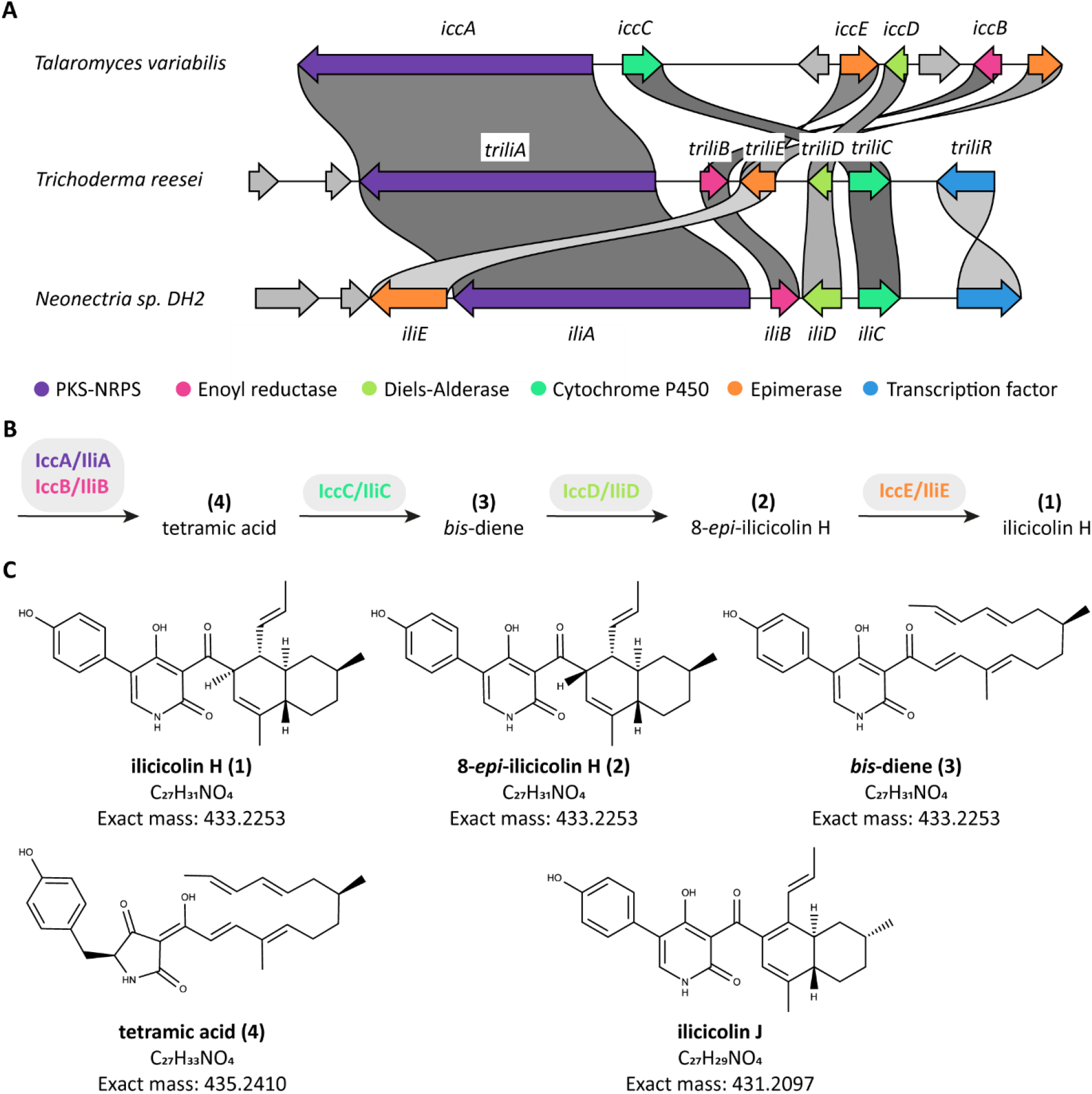
(A) The ilicicolin BGCs of *T. reesei* QM6a, *T. variabilis* HXQ-H-1, and *Neonectria* sp. DH2 were compared using the clinker tool^15^. The input .gbk files are provided in the supplemental materials. The *Neonectria* epimerase gene is comparatively larger, maybe due to a mistake of the gene model (prediction contains a TIM-like epimerase domain as well as a DnaJ-class molecular chaperone domain). (B) Model for the biosynthetic pathway for ilicicolin H, described by *Zhang* et al.^12^ and partially consistent with *Lin* et al.^13^ and *Shenouda* et al.^14^. (C) Previously described and for this publication relevant ilicicolin compounds; **(1)** – **(4)** from *Zhang* et al.^12^, ilicicolin J from *Lin* et al.^13^.

Known as a potent and broad-spectrum antifungal compound^2^ that specifically inhibits the cytochrome bc1 complex (reductase)^3^, the agent was also applied recently in several *in vitro* studies for treatment of various cancer cell lines (DLD-1 colon adenocarcinoma and A549 non-small cell lung carcinoma^4^, PC-3 and 22Rv1 prostate carcinoma^5^, Huh7 and HepG2 hepatocellular carcinoma^6^). Notably, ilicicolin H exhibits a manifold higher potency against fungi compared to mammalian cells, with half maximal inhibitory concentrations (IC_50_) of 2 – 3 ng mL^-1^ for *C. albicans* MY1055 NADH:cytochrome bc1 reductase compared to 2000 – 5000 ng mL^-1^ for rat liver cytochrome bc1 reductase^7^. Although ilicicolin H appeared as a promising treatment for yeast infections, it only showed modest efficacy in *in vivo* mouse models, which was attributed to high plasma protein binding^2^. Several attempts were undertaken to improve the efficacy by chemically modifying its structure, however, with limited success^2,7,8^.

Ilicicolin H is produced by several fungi during cultivation, e.g. *Cylindrocladium ilicicola* (strain MFC-870)^1,9,10^, *Gliocladium roseum*^2,11^, *Nectria* sp. B13^12^, and *Neonectria* sp. DH2^13^; and many additional fungi seem to possess the corresponding genes^12^. The biosynthetic gene cluster (BGC) of ilicicolin H was discovered by two groups independently in 2019. *Zhang* et al.^12^ and *Lin* et al.^13^ heterologously expressed the BGCs from *Talaromyces variabile* and *Neonectria* sp. DH2, respectively in *Aspergillus nidulans*. Recently, *Shenouda* et al.^14^ heterologously expressed the ilicicolin H BGC from *Trichoderma reesei* in *A. oryzae*. The BGCs of all three fungi contain similar genes with a polyketide synthase-nonribosomal peptide synthetase (PKS-NRPS) core gene (Figure 1A and Table 1).

**Table 1.** Proteins in the ilicicolin H BGC, comparison to the strains utilized in the heterologous experiments.

| Protein designation | Enzyme function | JGI Protein ID in *T. reesei* QM6a / <u>RutC30</u> | Homolog in *Neonectria* sp. DH2 (% identity) | Homolog in *T.s variabilis* HXQ-H-1 (% identity) |
| --- | --- | --- | --- | --- |
| TriliA | PKS-NRPS | 58285 / 128011 | IliA (63 %) | IccA (59.8 %) |
| TriliB | Enoyl reductase | 58289 / 74247 | IliB (65.3 %) | IccB (68.6 %) |
| TriliC | Cytochrome P450 | 58953 / 33667 | IliC (70.7 %) | IccC (69.4 %) |
| TriliD | Diels-Alderase | ----- / 75073 | IliD (55.6 %) | IccD (51.8 %) |
| TriliE | Epimerase | 76204 / 96728 | IliE (52 %) | IccE (55.9 %) |
| TriliR | Transcription factor | 72993 / 74475 |  |  |

Independent of the source organism, organization of the ilicicolin H BGC is in wide parts conserved, resulting in the following model (Figure 1B) for the biosynthetic pathway: first, the polyketide synthase (PKS) part of IccA/IliA assembles the polyketide backbone, supported by the enoyl reductase IccB/IliB, and adds the methyl groups. The nonribosomal peptide synthetase (NRPS) part of IccA/IliA then adds a tyrosine. Following a Dieckmann condensation, a tetramic acid intermediate **(4)** is released. This moiety is converted to the pyridone part of ilicicolin H by the cytochrome P450 IccC/IliC via a ring expansion. Next, the Diels-Alderase IccD/IliD is suggested to catalyze the intramolecular Diels-Alder reaction yielding the decalin moiety of ilicicolin H. Up to this step, the results of the three groups concur.

*Lin* et al.^13^ and *Shenouda* et al.^14^ report that these four enzymes from *Neonectria* sp. DH2 and *T. reesei*, respectively, are sufficient to obtain ilicicolin H in their heterologous expression experiments, whereas *Zhang* et al.^12^ demonstrate that IccE is necessary to catalyze the epimerization of 8-*epi*-ilicicolin H **(2)** to ilicicolin H **(1)** *in vivo* and *in vitro*. *Lin* et al.^13^ speculate that IccD and IliD differ, resulting in the formation of 8-*epi*-ilicicolin H and ilicicolin H, respectively. The epimerization appears to be pH-dependent^12^ and the observed epimerization differences might thus be a result of varying cultivation and/or extraction conditions. Further, the possibility that a host enzyme might also carry out the epimerization has to be considered. It is unclear whether *Zhang* et al.^12^ and *Lin* et al.^13^ used the same *A. nidulans* strain, since *Lin* et al.^13^ did not specify a strain designation. Importantly, there were other differences observed in the studies of these two groups that might be the result of different hosts. Most prominently, the heterologous expression conducted by *Lin et al.*^13^ did not only produce ilicicolin H but also a shunt product with comparable antifungal activities, ilicicolin J (Figure 1C). Similarly, *Shenouda et al.*^14^ reported on several acetylated side products during the heterologous expression of the *T. reesei* BGC in *A. oryzae* which they explained as a result of the native metabolism of the production host.

Herein, we describe the high-yield expression of ilicicolin H by genetic BGC activation in a native host for the first time. To this end, we overexpressed the BGC transcription factor, TriliR, under the constitutive *tef1* promotor^16^ in *T. reesei*. Further, we deleted the genes *triliA* and *triliE* to elucidate their roles in the ilicicolin H biosynthesis in the native host *T. reesei*. Overexpression of the cluster resulted in the production of three isobaric products (Figure 1C), namely ilicicolin H **(1)**, 8-*epi*-ilicicolin H **(2)** and the *bis*-diene **(3)**, and shed further light on the production process and the roles of the enzymes involved. Moreover, during this analysis, we uncovered a novel antifungal ilicicolin H derived compound, which we termed ilicicolin K.

## 2. Material and Methods

### 2.1. Material

All chemicals used in this study were sourced as follows: unless otherwise specified, all chemicals were purchased from Sigma-Aldrich (St. Louis, MO, USA). Phenol was purchased from AppliChem (Darmstadt, Germany). Malt extract, peptone, isoamylalkohol, FeSO_4_, CoCl_2_*6H_2_O, Na_2_MoO_4_*H_2_O, (NH_4_)SO_4_ and KH_2_PO_4_ were acquired from Merck (Darmstadt, Germany). Isopropanol, MgSO_4_*7H_2_O, CaCl_2_*2H_2_O, Na_2_HPO_4_*2H_2_O and uridine were obtained from Carl Roth (Karlsruhe, Germany). Agar and chloroform were purchased from VWR (Radnor, PA, USA). LC-MS grade acetonitrile (ACN) was acquired from VWR chemicals (Radnor, PA, USA). LC-MS grade methanol (MeOH) was obtained from Honeywell (Muskegon, MI, USA). LC-MS grade 2-propanol (IPA) was obtained from Fisher Scientific (Hampton, NH, USA). HPLC grade Diethyl ether was obtained from Sigma-Aldrich (St. Louis, MO, USA). Water (H_2_O) was purified in-house using a Barnstead™ Smart2Pure™ Water Purification System from Thermo Fisher Scientific (Waltham, MA, USA). Ilicicolin H (CAS #12689-26-8) standard, Catalog #10-3243 (Lot #X107453), was purchased from Focus Biomolecules (Plymouth Meeting, PA, USA).

Mycelium samples were lysed using the Bead Mill Max Homogenizer in combination with Tough Microorganism Lysing Mix Glass Beads (both VWR International, Radnor, PA, USA). Sonication was carried out using a Branson SFX550 sonifier from Emerson (Ferguson, MO, USA). Isolation of ilicicolin H from the medium was conducted using the Supelclean™ LC-18 SPE Tubes (Merck, Darmstadt, Germany).

Four biological replicates of fungal mycelium or culture supernatant were processed for each of the constructed and utilized strains, resulting in a total of four samples per experimental condition, unless otherwise noted.

### 2.2. Strains and cultivation conditions

All *T. reesei* strains (Table 2) were maintained on malt extract (MEX) plates (3 % malt extract, 0.1 % peptone, 1.5 % agar). 5 mM uridine or 25 µL per 100 mL Hygromycin B (Millipore, 400051) were added if required. For liquid cultivations 10^9^ spores L^-1^ were inoculated in Mandels Andreotti (MA) medium^17^ (KH_2_PO_4_ 2 g L^-1^, (NH_4_)_2_SO_4_ 1.4 g L^-1^, Urea 0.3 g L^-1^, FeSO_4_·7H_2_O 0.005 g L^-1^, MnSO_4_·H_2_O 0.0016 g L^-1^, ZnSO_4_·7H_2_O 0.0014 g L^-1^, CoCl_2_ 0.002 g L^-1^, MgSO_4_·7H_2_O 0.3 g L^-1^, CaCl_2_ 0.3 g L^-1^, peptone 0.75 g L^-1^, carbon source 10 g L^-1^) and incubated at 30 °C at 180 rpm.

**Table 2.** Utilized strains.

| Strain designation | Genotype | Source |
| --- | --- | --- |
| QM6a Δmus53, “wildtype” | Δmus53 | Steiger et al.<sup>18</sup> |
| QM6a Δpyr4 | Δmus53 Δpyr4 | Derntl et al.<sup>19</sup> |
| QM6a OETriliR | Δmus53 *Ptef::triliR::Tcbh2* (upstream of *pyr4*) | This study |
| QM6a ΔTriliA | ΔtriliA::hygR in QM6a OETriliR | This study |
| QM6a ΔTriliE | ΔtriliE::hygR in QM6a OETriliR | This study |

### 2.3. Genetic constructions

All PCRs were performed with the Q5 High-Fidelity DNA Polymerase (NEB) according to the manufacturer’s instructions. For cloning purposes, the *Escherichia coli* strain Top10 (Invitrogen) and the *Saccharomyces cerevisiae* strain WW-YH10 (ATCC 208405) were used. All plasmids and genetic constructs were verified by sequencing at Microsynth.

For the overexpression of TriliR the plasmid pRP4-OETriliR was constructed. First, a NotI-free coding region of *triliR* was constructed by a SOE-PCR using the primers TriliR_fwd-AflII, TriliR _MRev_SOE, TriliR _MFwd_SOE, and TriliR _rev-SpeI and chromosomal DNA of *T. reesei* QM6a Δmus53 as template yielding a coding region with a silent mutation at R227. This modified coding region was inserted into pRP4-TX(WT)^20^ via digestion with AflII and SpeI (both NEB) and ligation with T4 DNA Ligase (NEB). Approx. 20 µg of the plasmid were linearized with NotI, precipitated with sodium acetate and ethanol and dissolved in 15 µL ddH_2_O before the transformation into *T. reesei* (Figure S1).

For the construction of the *triliA* deletion cassette, a yeast recombinational cloning was performed using the lithium acetate method^21^ for the yeast transformation. The plasmid pRS426^22^ was linearized by digestion with KpnI and HindIII. The flanking regions were amplified by PCR with the primers 5_ TriliA _fwd-pRS426 and 5_ TriliA _fwd-pRS426 or 3_ TriliA _fwd-hph and 3_TriliA_rev-pRS426 using chromosomal DNA of *T. reesei* QM6a Δmus53 as template. The hph gene was amplified by PCR with the primers hph_fwd-5_ TriliA and hph_rev-3_ TriliA using the plasmid pAN7-1^23^ as template. The plasmid was extracted from yeast using the Zymoprep Yeast Plasmid Miniprep Kit (Zymo Research) and transformed into *E. coli* Top10 for amplification. To delete *triliA*, a split marker approach was used (Figure S2). To this end, a fusion of the 5’flank and a part of the hygR were amplified by PCR using the primers 5_ TriliA _fwd-pRS426 and hph_MR and the plasmid as template. Accordingly, the remaining part of the hph gene was amplified together the 3’flank using the primers hph_MF and 3_TriliA_rev-pRS426. Several PCR reactions were pooled, precipitated with ethanol and dissolved in 15 µL ddH_2_O.

For the deletion of *triliE*, also a split marker approach (Figure S3) was used, but the fusion products were directly constructed by SOE-PCRs. For the hph gene (from pAN7-1^23^) fragments, the primers PgpdA_fwd, hph_MR, hph_MF, and TtrpC_rev were used. The flanking regions were amplified by PCR with the primers triliE, the primers TriliE_5fwd, TriliE _5rev-hph, TriliE _3fwd-hph, and TriliE _3rev using chromosomal DNA of *T. reesei* QM6a Δmus53 as template. The fusion PCR products were cloned into pJET1.2 using the CloneJET PCR Cloning Kit (Thermo Scientific). Before transformation the fusion products were amplified by PCR, several PCR reactions were pooled, precipitated with ethanol and dissolved in 15 µL ddH2O.

### 2.4. Fungal transformation

*T. reesei* was transformed using a polyethylene glycol-mediated transformation protocol of protoplasts. Spores of the recipient strain were plated on sterile cellophane sheets laid on MEX plates at 30 °C overnight. The mycelium was scraped off and transferred into 15 mL Buffer A (1.2 M sorbitol, 100 mM KH_2_PO_4_, pH 5.6) containing 600 mg Vinotaste Pro (Novozymes) and 0.5 mg chitinase from *Streptomyces griseus* (Sigma-Aldrich C6137). This mixture was incubated in a sterile petri dish in an orbital shaker at 60 rpm and 30 °C for approx. 2-3 hours until the mycelium was completely disintegrated. The suspension was filtered through a 70 µm cell sieve and incubated on ice for 5 min. The suspension was filled up to 40 mL with ice-cold 1.2 M sorbitol and centrifuged at 2,500 g at 4 °C for 10 min. The pellet was resuspended in 30 mL 1.2 M sorbitol and again centrifuged. The protoplasts were finally resuspended in 1 mL ice-cold Buffer B (1 M sorbitol, 25 mM CaCl_2_, 10 mM Tris.Cl, pH 7.5). Next, the DNA (either 20 µg linearized plasmid or 5 µg of fusion PCR products each for the split marker approach) was filled up to 150 µL with ice-cold Buffer B, carefully mixed with 100 µL of the protoplast suspension and 100 µL “20 % PEG” (mixture of 6.7 mL Buffer B and 3.3 mL “60% PEG” (60 g PEG 4000, 1 mL 1 M Tris-HCl pH 7.5, 1 mL 1 M CaCl_2_, 38 mL ddH_2_O)) in a 50 mL reaction tube. This mixture was incubated on ice for 30 min before “60 % PEG” was added in steps (50 µL, 200 µL, 500 µL). Next, the tube was incubated at room temperature for 20 min, and finally, Buffer C was added in steps (200 µL, 400 µL, 1 mL, 2.5 mL). For plating, the tube was filled up to 50 mL with molten, 50 °C-warm selection medium (containing 1 M sucrose) and poured into a 14.5 cm petri dish. For insertion of the TriliR overexpression construct together with *pyr4* a minimal medium was used (MA medium with glucose without peptone, pH 5.8). For the deletion of *triliA* and *triliE* by replacement with the *hygR*, MEX medium with hygromycin was used. The plates were incubated at 30 °C under light until colonies were visible (up to a week). The candidates were then homokaryon selected by spore streaking on selection plates.

### 2.5. DNA extraction and genotyping

Mycelium was harvested and pressed dry between two sheets of filter paper. Approx. 50 mg were lysed in 1 mL CTAB buffer (1.4 M NaCl, 100 mM Tris-HCl pH 8.0, 10 mM EDTA, 2 % CTAB, 1 % polyvinylpyrrolidone) with 0.37 g small glass beads, 0.25 g medium glass beads and one large glass bead in a 2 mL screw cap reaction tube using a Fast-Prep-24 (MP Biomedicals, Santa Ana/, CA, USA) at 6 m s^-1^ for 30 sec. The samples were incubated at 65 °C for 20 min and finally centrifuged at 12,000 g for 10 min. The supernatant was transferred to a 2 mL reaction tube and the DNA was purified by a phenol-chloroform-isoamyl alcohol extraction, followed by two chloroform extractions. The samples were then treated with RNase A (Thermo Fisher Scientific) according to the manufacturer’s instructions and the DNA finally precipitated using isopropanol and dissolved in 10 mM Tris-HCl pH 8.0.

All PCR reactions for genotyping were performed with the OneTaq DNA Polymerase (NEB) according to the manufacturer’s instructions.

### 2.6. RNA extraction and RT-qPCR analyses

Mycelium was harvested and pressed dry between two sheets of filter paper, frozen in liquid nitrogen, and stored at −80 °C for up to a week. Approx. 50 mg of mycelium were disrupted in 1 mL RNAzol RT with 0.37 g small glass beads, 0.25 g medium glass beads and one large glass bead in a 2 mL screw cap reaction tube using a Fast-Prep-24 (MP Biomedicals) at 6 m s^-1^ for 30 sec. The samples were centrifuged at 12,000 g for 10 min, the supernatant was transferred to a 1.5 mL reaction tube and mixed with ethanol 1:1. The RNA was purified using the Direct-zol RNA MiniPrep Kit (Zymoreasearch) according to the manufacturer’s instructions. Notably, this kit contains a DNase treatment step. The total RNA was reverse transcribed using the LunaScript RT SuperMix Kit (NEB) according to the manufacturer’s instructions. The cDNA was diluted 1:50 in ddH_2_O and 2 µL used as a template in a 15 µL reaction using the Luna Universal qPCR Master Mix (NEB) on a Rotor-Gene Q (Qiagen). Primers were added and PCR reaction conditions were chosen according to the manufacturer’s instructions. To calculate the relative transcript abundance, we used the Pfaffl method^24^ and the *act1* and *sar1* genes as reference genes^25^.

### 2.7. Sample preparation for HPLC(-MS/MS) and NMR analysis

#### 2.7.1. Ilicicolin H standard

100 µg of ilicicolin H standard (Focus Biomolecules) were dissolved in 100 µL of DMSO for a final concentration of 1 µg µL^-1^. 1 µL of this stock was diluted with 99 µL of 50 % ACN, to obtain a final concentration of 0.01 µg µL^-1^. For LC-MS analysis, 1 µL was injected.

#### 2.7.2. Polar extracts of mycelium

100 mg of frozen mycelium sample were weighed into glass bead-milling tubes and 1 mL of polar extract composed of a mixture of acetonitrile/methanol/water (40:40:20) were added. Lysis was conducted by bead-milling (4x 30 sec, 6 m s^-1^) and subsequent sonication (30 s, 10 % intensity). The lysed samples were centrifuged for 10 min at 20,000 g and 20 °C and the supernatant was transferred into fresh Eppendorf tubes. The supernatants were centrifuged again, for 1 min at 20,000 g. For analysis, 10 µL of the supernatant were taken and diluted in 90 µL of 50 % ACN and 1 µL was injected for LC-MS analysis.

#### 2.7.3. Quantification of ilicicolin H in medium

For quantification of ilicicolin H in the media, a matrix-matched external calibration curve was utilized. For that, 4 mL of culture medium of each wildtype and ΔTriliA quadruplicate (n = 8 in total) were pooled, which do not contain ilicicolin H in detectable quantities (below LOD of 21.4 ng mL^-1^, Figure S5). 6 x 2 mL were taken out and each spiked with precise amounts of ilicicolin H standard (50 ng, 100 ng, 200 ng, 500 ng, 1000 ng and 2000 ng). These standards and the culture supernatants of the other 8 samples (4 x OETriliR, 4 x ΔTriliE, 50 mL each) were cleaned up using C18-SPE columns. The elution buffer was constituted of 10 % acetonitrile in isopropanol supplemented with 15 mM ammonium formate. The eluates were centrifuged for 10 min at 20,000 g before vacuum-drying them and reconstitution in 100 µL 50 % ACN each. 1 µL was used for LC-MS analysis.

#### 2.7.4. Proteomics

Proteomics analysis was conducted starting with 100 mg of frozen mycelium sample which were weighed into glass bead-milling tubes and 1 mL of reducing and alkylating buffer (100 mM TRIS HCl; pH = 8.5, 1 % sodium dodecyl sulphate, 10 mM tris(2-carboxyethyl) phosphine, 40 mM 2-chloroacetamide) was added. Lysis was conducted by bead-milling (2 min, 6 m s^-1^) and subsequent sonication (10 s, 10 % intensity). The lysed samples were spun down for 12 min at 20,000 g and 20 °C and the supernatant was transferred into fresh Eppendorf tubes. The supernatants were heated to 95 °C for 10 min at 330 rpm to perform reduction and alkylation of the proteins. 100 μg of protein per sample (according to bicinchoninic acid assay protein estimation (BCA), Thermo Fisher Scientific, reducing agent compatible) were subjected to acetone precipitation by adding NaCl to a final concentration of 10 mM and incubation for 5 min at room temperature. Subsequently, 4 x volumes of acetone were added and samples were incubated for 2 min. After centrifugation (15 min at 15,000 g) the supernatant was removed. Dried samples were dissolved in 25 % trifluoroethanol in 100 mM Tris-HCl (pH = 8.5) and subjected to vortexing and sonication until completely dissolved. For protein digest, samples were diluted to 10 % trifluoroethanol using 100 mM ammonium bicarbonate. Trypsin (Promega, Fitchburg, WI) was added in a 1:50 enzyme to protein ratio and digest was performed overnight at 37 °C and 500 rpm. The following day, 500 ng of digested sample were loaded on Evotips, according to the protocol of the manufacturer (Evosep, Odense, DK).

#### 2.7.5. Scaled up extraction for NMR analysis

Around 5 g of mycelium of both OETriliR and ΔTriliE strains were extracted by first grinding in liquid nitrogen to a fine powder and subsequently bead-milled in diethyl ether. After removing cell debris and beads, the diethyl ether extracts were washed two times with ddH_2_O, dried under a nitrogen stream, and reconstituted in 60 % ACN in 15 mM ammonium formate for subsequent HPLC separation and fractionation.

### 2.8. HPLC(-MS/MS) analysis

#### 2.8.1. Polar extracts of mycelium

After 1:10 dilution of the polar extracts, ilicicolin H (C_27_H_31_NO_4_) was measured employing an untargeted lipidomics workflow in positive polarization mode on a Bruker timsTOF Pro equipped with a VIP-HESI source (Bruker Corporation, Billerica, MA, USA). The frontend was a Thermo Fisher Scientific Vanquish H UHPLC with a Waters Acquity BEH C18 column (150 mm × 1 mm ID, 1.7 μm; Waters Corporation, Milford, MA, USA). Mobile phase A was 60 % acetonitrile in 15 mM ammonium formate; mobile phase B was 10 % acetonitrile in isopropanol supplemented with 15 mM ammonium formate. The following gradient was employed at 50 °C: 0 min; 2 % B; 100 µL min^-1^, 15 min; 100 % B; 70 µL min^-1^, 22 min; 100 % B; 70 µL min^-1^, followed by 5 min re-equilibration to starting conditions (2 % B; 100 µL min^-1^). The timsTOF Pro mass spectrometer (Bruker Daltonics, Germany) was operated in positive ionization mode with enabled trapped ion mobility spectrometry (TIMS) at 100 % duty cycle (100 ms ramp time). Source capillary voltage was set to 4500 V and dry gas flow to 8 L min^-1^ at 230 °C. Sheath Gas Flow was set to 4.0 L min^-1^ at a temperature of 100 °C, with active exhaust being activated. Scan mode was set to parallel accumulation serial fragmentation (PASEF) with a scan range from 300 to 1300 *m/z*, using a mobility (1/K0) window from 0.8 to 1.81 V*s cm^-2^. Collision Energy was set to 45.00 eV. Full MS method details are available as metadata in the raw data file deposited in the Pride repository.

To obtain fragmentation spectra for all features without pre-filtering by ion mobility, measurements for molecular networking were conducted without trapped ion mobility spectrometry (TIMS off). The LC eluents and gradient remained unaltered. The timsTOF Pro mass spectrometer (Bruker Daltonics, Germany) was operated in positive ionization mode, with the same settings as described above. Only the Sheath Gas Flow temperature was changed to 200 °C. Scan mode was set to Auto MS/MS with 12.00 Hz MS spectra rate and 16.00 Hz MS/MS spectra rate, resulting in a total cycle time of 0.5 s. Scan range was set from 20 to 1300 *m/z*.

#### 2.8.2. Quantification of ilicicolin H in medium

HPLC-MS/MS measurement of the prepared standards and extracts were precisely carried out as described in chapter 2.8.1.

#### 2.8.3. Proteomics

500 ng of the digest were separated on the Evosep One equipped with an Ionopticks Aurora Series UHPLC C18 column (15 cm x 75 µm ID, 1.7 µm; Ionopticks, Fitzroy, VIC, Australia). The LC–method Whisper_40SPD was used with solvent A being 0.1 % formic acid in water and solvent B acetonitrile containing 0.1 % formic acid while maintaining the column at 40 °C. The timsTOF HT mass spectrometer (Bruker Daltonics, Germany) was operated in positive mode with enabled trapped ion mobility spectrometry (TIMS) at 100 % duty cycle (100 ms ramp time). Source capillary voltage was set to 1500 V and dry gas flow to 3 L min^-1^ at 180 °C. Scan mode was set to data independent parallel accumulation–serial fragmentation (diaPASEF) using parameters previously optimized with py_diAID^26^. In brief, 24 isolation windows from *m/z* 300 to 1,200 and 1/K0 0.7 to 1.35 were defined. After MS1 scan, 2 isolation windows were fragmented per TIMS ramp resulting in an overall DIA cycle time of 1.38 s. Full MS method details are available as metadata in the raw data file deposited in the Pride repository.

#### 2.8.4. Scaled up extraction for NMR analysis

Separation and fractionation of the mycelial diethyl ether extracts was conducted on a Thermo Fisher Scientific Vanquish F UHPLC with a Waters XBridge BEH C18 column (150 mm × 4.6 mm ID, 2.5 μm; Waters Corporation, Milford, MA, USA). Mobile phase A was 60 % acetonitrile in 15 mM ammonium formate; mobile phase B was 10 % acetonitrile in isopropanol supplemented with 15 mM ammonium formate. The following gradient was employed at 50 °C and a flow-rate of 1 mL min^-1^, for OETriliR samples: 0 min; 2 % B, 10 min; 45 % B, 11 min; 98 % B; 15 min; 98 % B, followed by 5 min re-equilibration to starting conditions (2% B; 1 mL min^-1^). UV-absorption was detected at a wavelength of λ = 320 nm. For ΔTriliE samples, the gradient was slightly adapted for better isomer separation: 0 min; 2 % B, 7 min; 33.5 % B, 10 min; 40 % B; 11 min; 98 % B, 15 min; 98 % B, followed by 5 min re-equilibration to starting conditions. Flow-rate and separation temperature remained unchanged (1 mL min^-1^, 50 °C). Fractions were manually collected, united, and dried under vacuum. Dried extracts were quantified gravimetrically and 6.73 mg ilicicolin H were extracted from the 5 g dried OETriliR mycelium employed for NMR analysis.

To purify ilicicolin H and K for the microbial tests, diethyl ether extracts from OETriliR mycelia were fractionated on an Autopurification system of Waters using an ACQUITY QDa Detector in combination with a 2998 Photodiode Array Detector, equipped with a XSELECT CSH C18 OBD Prep column (150 mm × 30 mm ID, 5 μm; Waters Corporation, Milford, MA, USA). The following gradient was employed at room temperature (25 °C) and a flow-rate of 20 mL min^-1^: 1 min; 2 % B, 21 min; 45 % B, 23.40 min; 98 % B; 36 min; 98 % B, 37 min; 2 % B, followed by 13 min re-equilibration to starting conditions (2 % B; 20 mL min^-1^). The PDA detector was set to a wavelength range from λ = 200 nm to λ = 450 nm, with a resolution of 1.2 nm and a sampling rate of 10 points s^-1^.

### 2.9. NMR analysis

All samples were measured on a Bruker Avance III 600 MHz spectrometer with Prodigy nitrogen cryo BBFO{H-F} inverse probe head. Spectra were recorded at 298 K and are referenced to the residual solvent signal.

### 2.10. Antifungal activity assays

To determine the broth dilution minimum inhibitory concentration (MIC) of the isolated substances, we performed microdilution tests according to the EUCAST guidelines (https://www.eucast.org/astoffungi/methodsinantifungalsusceptibilitytesting; EUCAST E.Def 7.4 October 2023 and EUCAST E.DEF 9.4 March 2022). In brief, ilicicolin H and ilicicolin K were dissolved in DMSO to a concentration of 25.6 mg mL^-1^. These stock solutions were serially diluted 1:2 to obtain the working solutions 12.8, 6.4, 3.2, 1.6, 0.8, 0.4, 0.2, 0.1, and 0.05 mg mL^-1^. All solutions were added 1:100 to double strength RPMI 2 % G supplemented with Tween-20 (RPMI 1640 with L-glutamine without Sodium Bicarbonate (Thermo Scientific, Catalog number: 31800089), 20.8 g L^-1^; MOPS, 69.06 g L^-1^; glucose, 36 g L^-1^; Tween-20, 0.004 % (v/v); pH adjusted to 7.0 with NaOH). For the *S. cerevisiae* cultivations, uracil was added to a concentration of 40 mg mL^-1^ to the double strength RPMI 1640 2 % G. DMSO was added 1:100 to RPMI 2 % G to obtain a 0.0 mg mL^-1^ control. To prepare the *S. cerevisiae* inoculum, the strain CEN.PK113-5D^27^ was incubated on YPD plates (yeast extract, 10 g L^-1^; peptone, 20 g L^-1^; glucose, 20 g L^-1^) at 30 °C for 48 hours until individual colonies were obtained. Several colonies were suspended in sterile distilled water to a density of 0.5 McFarland. To the *A. nidulans* inoculum the strain FGSC A4 (CBS 112.46) was incubated on potato dextrose agar (PDA) plates at 30 °C for a week. Spores were harvested by rolling a sterile cotton swab on the colony and resuspending them in sterile distilled water. The spore suspension was filtered through glass wool and adjusted to a density of 0.5 McFarland. Finally, 100 µL of the supplemented media were mixed with 100 µL of inoculums in a well of a sterile, 96-well microdilution plate with flat-bottom wells, resulting in the test concentrations 128, 64, 32, 16, 8, 4, 2, 1, 0.5, 0.25 µg mL^-1^ of ilicicolin H and K. The assay was performed in technical triplicates, and the medium was mixed with sterile water instead of the inoculums used as blank. The well plates were incubated at 30 °C and the optical density at 560 nm was measured using a plate reader (Promega ProMax). We defined a growth inhibition of at least 50 % in comparison to the growth in the DMSO-supplemented medium as the cut-off to determine the MIC.

### 2.11. Data analysis

#### 2.11.1. General utilized software

MarvinSketch 22.6.0 (Chemaxon, www.chemaxon.com) was used for drawing, displaying and characterizing chemical structures, substructures and reactions. For fragmentation spectra analysis, SIRIUS version 5.8.6 was used: SIRIUS^28^ for molecular formula identification and CSI:FingerID^29,30^ for structure database search and substructure annotation. MZmine version 4.0.3 (https://mzio.io/) was used for feature finding and to conduct molecular networking^31,32,33^.

#### 2.11.2. Polar Extracts

Ilicicolin H (C_27_H_31_NO_4_) was quantified on MS1 level employing the open-source application Skyline version 22.2^34^. Mass-to-charge, retention time, and CCS were manually matched to an authentic standard.

#### 2.11.3. Quantification of ilicicolin H in medium

Ilicicolin H (C_27_H_31_NO_4_) was quantified in standards and samples on MS1 level employing the open-source application Skyline version 22.2^34^. As usual, mass-to-charge, retention time, and CCS were manually matched to an authentic standard. Using the areas and known concentrations of the matrix matched standards, an external calibration curve was created (Figure S6).

#### 2.11.4. Proteomics

Proteomics data was analyzed using the software DIA-NN 1.8.1^35,36^. For *Trichoderma reesei (Hypocrea jecorina*, taxonomy ID 51453, 19,246 entries), all reviewed (Swiss-Prot) and unreviewed (trEMBL) FASTA files were downloaded from UniProtKB (https://www.uniprot.org/) on the 26^th^ of June 2023. The database was manually extended with common contaminants (https://www.thegpm.org/crap/) resulting in 19,366 entries, and used for a library-free search with FDR set to 1 %. Deep learning-based spectra, retention time and ion mobility prediction were enabled, minimum fragment ion m/z was set to 200 and maximal ion fragment m/z to 1800. Trypsin was employed as protease and the maximum number of missed cleavages was configured with 2. Peptide length was adjusted to a range from 7 to 30 amino acids. Cysteine carbamidomethylation was set as a fixed and methionine oxidation as a variable modification, allowing a maximum of one variable modification. DIA-NN optimized the mass accuracy automatically using the first run in the experiment.

This resulted in a list of 6,039 proteins with their corresponding LFQ values. Data processing using protein group quantities was done with Perseus software version 2.0.11^37^. Intensities were log2 transformed and contaminants removed. The matrix was then filtered to contain 100 % valid values in at least one group. This reduced the matrix to 5,977 proteins, and remaining missing values were imputed from normal distribution (downshift 1.8, width 0.3). Histograms of data distribution per each sample highlighting imputed values is displayed in Figure S7. Principal component analysis was performed without category enrichment. Volcano blots of protein expression differences were generated using the following criteria: p-value of 0.05, S0 of 0.1 and permutation-based FDR set to 5 % to correct for multi-testing with 250 randomizations.

## 3. Results

### 3.1. Activation of the ilicicolin H BGC by overexpression of TriliR

The ilicicolin H BGC contains, next to the biosynthetic genes *trIliA-E*, also a gene encoding for a transcription factor (Protein ID 72993, Figure 1A). The JGI gene model^38^ suggests a coding region with 1,389 bp encoding for a protein with 462 aa (Supplemental File TriliR_gene_models.fasta) containing only a partial fungal transcription factor middle homology region (FTFMHR, cd12148 of the conserved domain database^39^) at residues 16-231 with an E-value of 4.82e-15 but no DNA-binding domain^40^. An alternative gene model (Protein ID 74475) is suggested for *T. reesei* strain RutC-30^41^ (Supplemental File TriliR_gene_models.fasta); this model does not contain any introns, consists of 2,364 bp encoding for a protein with 787 aa, and it contains a full-length FTFMHR at residues 270-733 (E-value 1.34e-27) and a GAL4-like zinc cluster DNA binding domain at residues 11-47 (E-value 9.28e-10). Consequently, we used the RutC-30 gene model for our molecular biological work and refer to the gene and protein as *triliR* and TriliR, respectively.

To overexpress TriliR, we put an altered genomic sequence (silent mutation at R227, Supplemental File TriliR_gene_models.fasta) under the control of the constitutive promoter of *tef1* and inserted it in front of the *pyr4* locus into the *T. reesei* strain QM6a Δpyr4 as previously described^19^ (Figure S1). We could detect substantially higher transcript levels of all biosynthetic genes in the ilicicolin H BGC in the strain OETriliR (approx. 300 to 1,500 times higher), compared to the control strain (Supplemental File RT-qPCR.xlsx, Figure S4).

### 3.2. Proteomics & targeted ilicicolin H analysis

This successful overexpression on mRNA level was also confirmed on proteomic level, with all five enzymes TriliA – E being significantly upregulated in the OETriliR strain compared to the wildtype strain, as depicted in Figure 2A. Principal component analysis (PCA, Figure 2B) revealed that the conducted genetic modifications resulted in the emergence of three distinct clusters. Specifically, sole overexpression of the ilicicolin H cluster (strain designation: OETriliR) leads to a marked change in the protein expression profile, evident from its clustering distant to the wildtype samples. Samples with ilicicolin-BGC overexpression and simultaneous knockout of the epimerase *triliE* (strain designation: ΔTriliE), the enzyme catalyzing the last step of ilicicolin H formation, demonstrate an overlap with the OETriliR cluster, suggesting minimal proteomic changes following the *triliE* knockout. On the contrary, upon ilicicolin-BGC overexpression and simultaneous knockout of *triliA* (strain designation: ΔTriliA), which initiates ilicicolin H formation, a more pronounced proteomic change is induced, apparent by clustering apart from the others yet closer to the wildtype.

**Figure 2.**
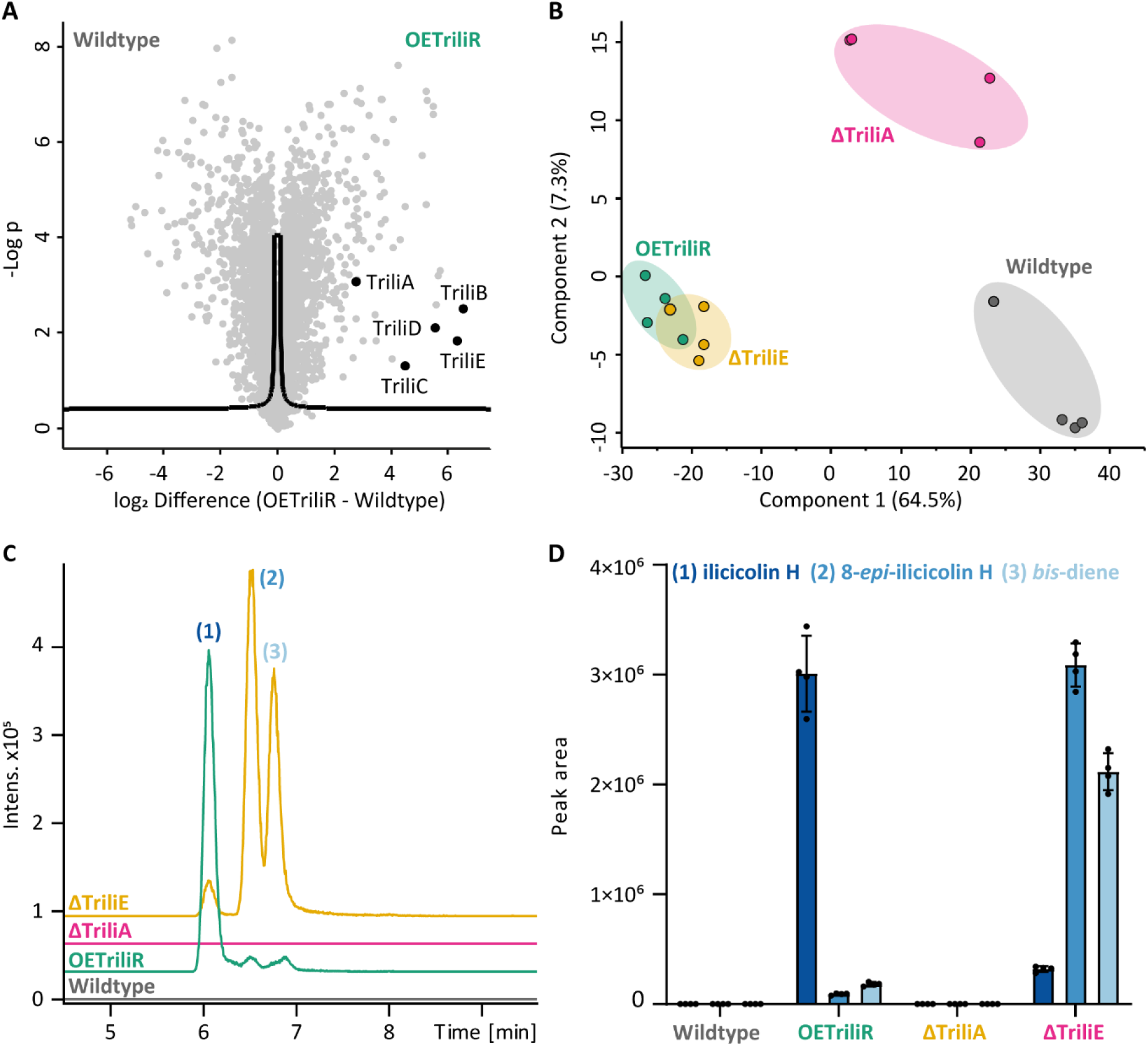
(A) Significant upregulation of all five enzymes involved in ilicicolin H production on protein-level, OE vs. WT (FDR: 0.05, s: 0.1). (B) PCA clustering of the four strains. (C) Extracted ion current chromatograms (EICCs) for ilicicolin H (C27H31NO4, [M+H]^+^ 434.2326 ± 0.02) of the four utilized strains. (D) MS1 Peak areas of ilicicolin compounds (n = 4).

Targeted metabolomics analysis of ilicicolin H revealed that the *T. reesei* wildtype strain does not produce ilicicolin H or direct biochemical predecessors in quantities detectable by our assay (LOD of ilicicolin H: 21.4 ng mL^-1^, Figure S5) under basal conditions, indicating that the ilicicolin H biosynthetic gene cluster (BGC) is indeed silent under common laboratory conditions (Figure S4). Upon overexpression of TriliR (OETriliR), the BGC is successfully activated and ilicicolin H gets produced in high yields, as seen in the extracted ion current chromatograms (EICCs) in Figure 2C and D. Overall, more product can be found in the mycelium than in the medium, with values of 1.346 µg ilicicolin H per mg mycelium (**chapter 2.8.4**) vs. around 6 ng of ilicicolin H in the culture supernatant per mg mycelium (Table S3, factor ∼225 less; 50 mL medium). Upon knockout of *triliA* (ΔTriliA), the enzyme initiating ilicicolin H biosynthesis, the production of ilicicolin H is completely halted, as depicted in Figure 2C. This confirms that TriliA is necessary to initiate the formation process.

### 3.3. Ilicicolin H isomers in detail

Extracted ion current chromatograms of ilicicolin H reveal three isomers at the expected m/z (Figure 2C), with peak **(1)** representing the main product, whereas peak **(2)** represents 8-*epi*-ilicicolin H and peak **(3)** the *bis*-diene, respectively. The epimerase TriliE catalyzes the last step in the ilicicolin H biosynthesis, namely the epimerization from 8-*epi*-ilicicolin H **(2)** to ilicicolin H **(1)**^12^. **(2)** in turn is a product of a putative S-adenosylmethionine (SAM)-dependent Diels-Alderase^14^, that catalyzes the transformation from *bis*-diene **(3)** to 8-*epi*-ilicicolin H **(2)**. Knockout of the epimerase encoding gene *triliE* leads to a strong decrease in the production of **(1)**, and instead, **(2)** and **(3)** increase drastically in their abundance (Figure 2C and D). This is in contrast to the previous report by *Zhang et al*.^12^, where deletion of the epimerase-encoding gene yielded mainly 8-*epi*-ilicicolin H **(2)** and the biosynthetic precursor of the *bis*-diene, the tetramic acid **(4)**. A possible explanation for that could be that the ring-expanding cytochrome P450 (IccC^12^, TriliC^14^), responsible for the conversion of the initially formed tetramic acid to the *bis*-diene^12^, is more efficient in the native *Trichoderma reesei* host system. This might be either be caused by higher expression in the engineered BGC or, more likely, by a better cofactor supply by a matched P450 reductase^42^.

The fact that ilicicolin H is still present even if *triliE* is deleted, albeit at a significantly lower level, corroborates the results by *Zhang* et al.^12^, who showed that nonenzymatic epimerization of **(2)** to **(1)** can occur, more preferably at higher pH values. However, in the presence of TriliE the epimerization process is now remarkably improved, underlining the necessity of this enzyme to produce the final product ilicicolin H in high yields in *T. reesei*.

### 3.4. Molecular networking and discovery of the novel compound ilicicolin K

Molecular networking of the untargeted metabolomics data using MZmine 4 combined several features to an “ilicicolin” network (Figure 3). In addition to ilicicolin H **(1)** and its pathway intermediates **(2)** and **(3)**, a fourth compound with a monoisotopic mass of 433.2252 Da was detected. This compound, like **(1)**, **(2)** and **(3)**, exhibited the same ionization adducts ([M+H]^+^, [M+H-H2O]^+^, [M+Na]^+^) but eluted substantially later than the other three. Despite similar fragmentation spectra and *m/z* ratio, this feature is unlikely to be ilicicolin H or one of its measured intermediate products because of a vastly different retention time (RT = 8.23 min vs 5.84 min for **(1)**, 6.31 min for **(2)** and 6.57 min for **(3)**) and the feature not being visible when ion mobility is turned on, in the applied mobility (1/K0) window from 0.8 to 1.81 V*s cm^-2^. To the network nodes of **(1)** and **(2)**, five additional features were clustered with *m/z* values of 432.2169, 450.2273, and 448.2117 (feature IDs 3, 5, 6, 9, 16 and 21). Another feature with *m/z* 448.2114 (ID 9) was matched to both *m/z* 448.2114 nodes (IDs 5 and 6). Manual inspection revealed that this feature was an artifact of the feature detection of the ilicicolin H **(1)** main peak, rendering it redundant.

**Figure 3.**
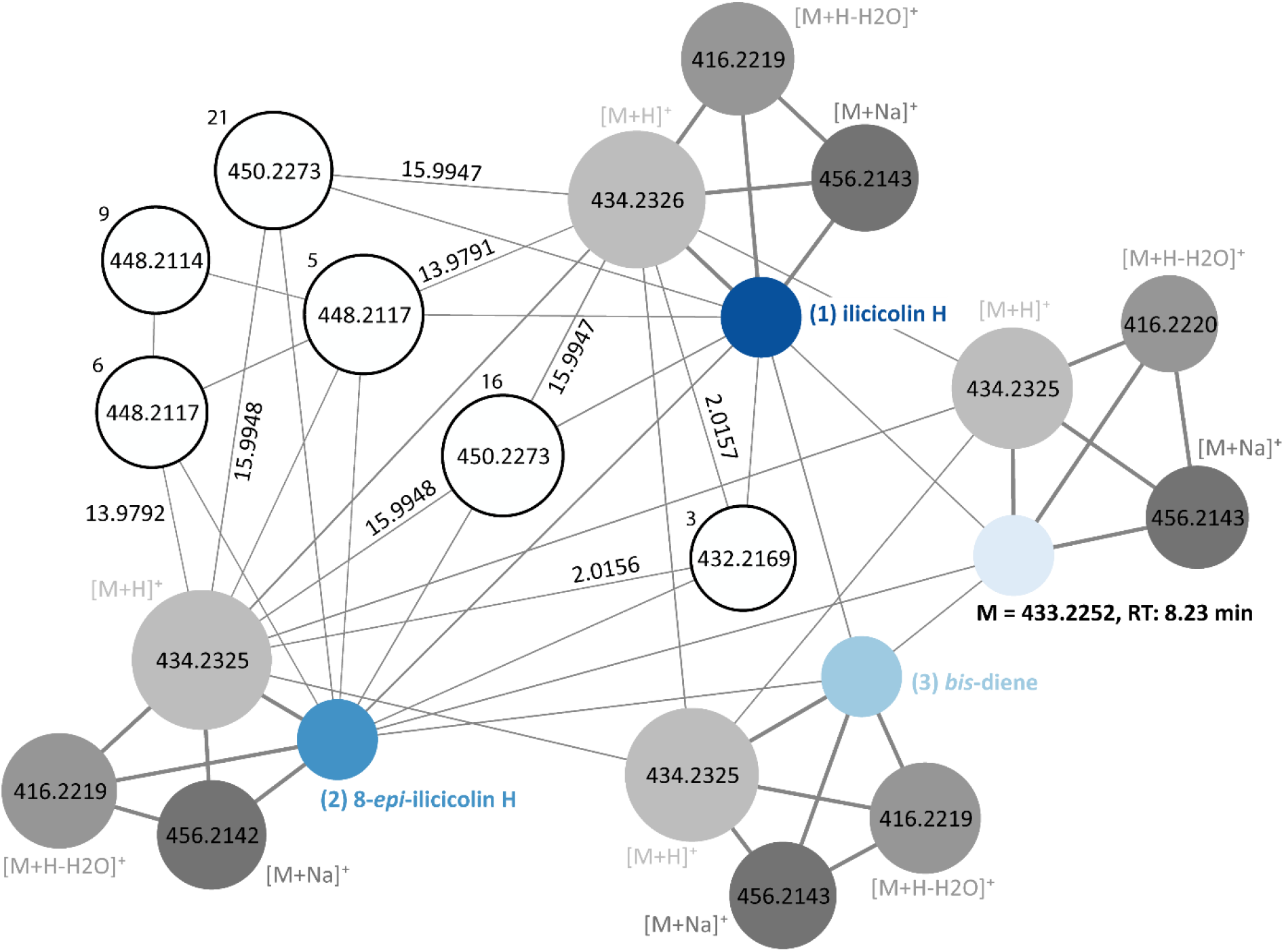
Ion-identity molecular networking using MZmine 4 shows the connections between the previously known ilicicolin H compounds **(1)**, **(2)** and **(3)** and reveals the presence of several related compounds. Size of the nodes corresponds to the log10_sum_intensity of the features, width of the edges to the similarity score. Grey colors indicate ionization adducts ([+H], [+Na], [+H-H2O]). White nodes with black contour represent unknown features, the numbers next to them indicate their feature ID within the network.

Molecular formula prediction by SIRIUS revealed for *m/z* 432.2169 the sum formula C_27_H_29_NO_4_, which could correspond to the compound ilicicolin J (Figure 1C). For *m/z* 450.2273, C_27_H_31_NO_5_ was proposed, implying an oxidized ilicicolin H variant. For *m/z* 448.2117, C_27_H_29_NO_5_ was suggested, which corresponds to a neutral mass of 447.2040 Da. This unusual modification - an oxidation with simultaneous loss of two hydrogen atoms - caught our attention and we conducted further investigation by comparing its fragment spectra to those of **(1)**, **(2)** and **(3)** in Figure 4.

**Figure 4.**
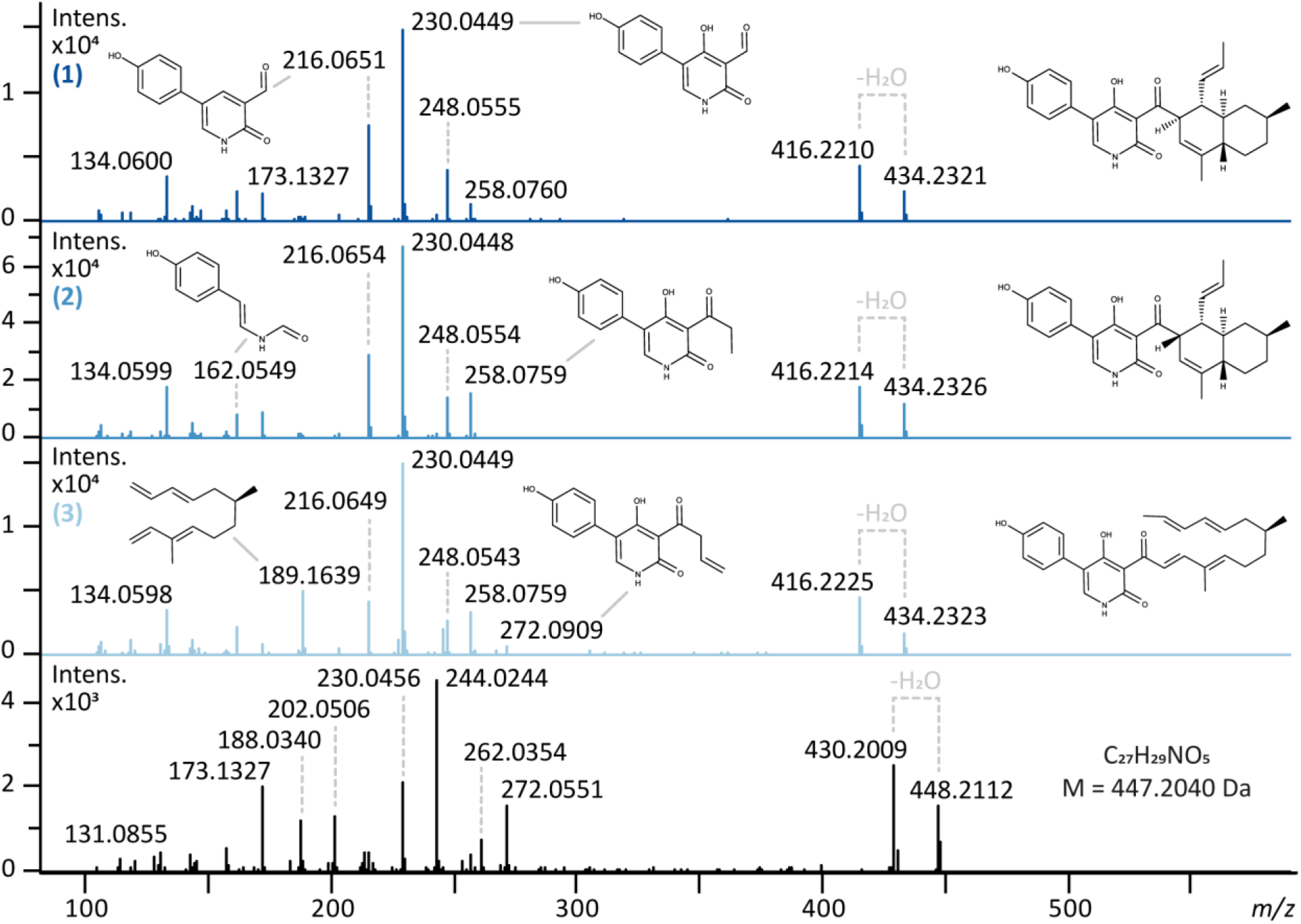
Fragmentation spectra of ilicicolin H **(1)**, 8-*epi*-ilicicolin H **(2)**, the *bis*-diene **(3)** and the novel ilicicolin compound with the sum formula C27H29NO5. Substructure annotation was conducted using SIRIUS.

Although **(1)**, **(2)** and **(3)** are isomeric compounds with the sum formula C_27_H_31_NO_4_ and an exact mass of 433.2253 Da, their fragmentation spectra show certain differences (Figure 4). Whereas **(1)** and **(2)**, the two epimers, reveal nearly identical fragments with very similar intensities, **(3)** differs in regard to fragments which are part of the open chain (decalin unit) of the molecule. This becomes apparent by an intense fragment with *m/z* 189.1639, which is far less pronounced in **(1)** and **(2)**, and the fragment *m/z* 272.0909. The first one constitutes the fragmented decalin unit, which fragments more readily in open conformation than when having undergone the Diels-Alder cycloaddition. The second one is the phenyl-pyridone unit with a part of the decalin unit attached. Overall, all three compounds fragment preferably directly between these two moieties, resulting in the most pronounced ion with *m/z* 230.0449. The novel compound shows a characteristic shift of + 13.98 Da for many of the fragments, compared to the spectrum of **(1)**. Judging from the location of the shift, the modification is most likely located in the phenyl-pyridone moiety of the molecule.

In accordance with these findings, NMR analysis revealed that the compound is actually an ilicicolin H derivative which was hydroxylated at the tyrosine moiety of the molecule with simultaneous ring formation of the hydroxyl group at the pyridone towards the tyrosine, yielding a dihydroxy-dihydro-oxa-azafluorene (Figure 5). We termed this novel compound ilicicolin K.

**Figure 5.**
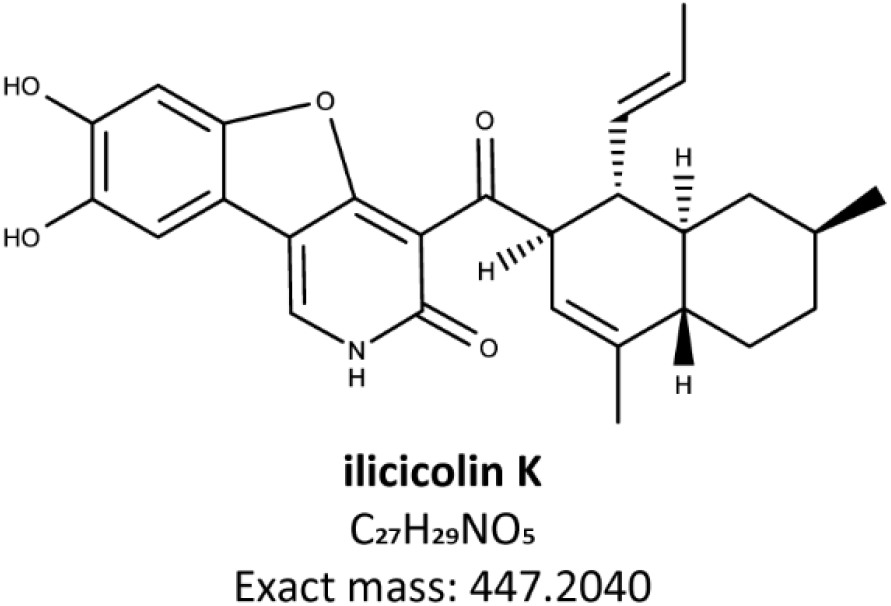
The newly discovered compound ilicicolin K.

Because of the structural similarities, we next tested whether ilicicolin K shares the antifungal characteristics of its biosynthetic relatives. Like ilicicolin H, ilicicolin K exhibits a strong antifungal activity on the tested strains *Saccharomyces cerevisiae* and *Aspergillus nidulans*. This is plausible since the β-diketone group of the molecule, which is essential for its activity^7^, remained unaltered compared to **(1)**. In order to determine the IC_50_ and MIC values against a variety of pathogenic fungal strains, further antimycotic and bioavailability tests are planned.

## 4. Discussion

Although the ilicicolin H biosynthetic gene cluster (BGC) has been previously described, earlier studies exclusively relied on heterologous expression systems to elucidate its pathway. In this study, we present the first successful genetic activation of the typically silent, albeit functional, ilicicolin H BGC of *T. reesei*, in its native host. We achieved this activation by overexpressing the cluster’s transcription factor, TriliR, under the control of the constitutive *tef1* promoter. Our results demonstrate successful expression of the cluster’s enzymes at both mRNA and protein levels. Furthermore, targeted metabolomics analysis confirmed high-yield production of ilicicolin H upon overexpression of TriliR.

In our system, the transcription factor TriliR is overexpressed, ensuring that the resulting expression levels of the enzymes TriliA – TriliE maintain their natural ratios relative to each other, mirroring normal physiological activation conditions. This approach allows for a precise assessment of each enzyme’s role within the cluster. In contrast, investigation of enzymatic functions in heterologous expression systems poses the risk of producing biased results because host enzymes can promote or interfere with the artificial pathway, potentially leading to the formation of undesired compounds or shunt products that would not naturally occur in the native host.

This potential discrepancy becomes apparent when comparing our findings to those of *Shenouda et al.*^14^ who investigated the same BGC from *T. reesei* through heterologous expression in *A. oryzae.* The authors registered high amounts of ilicicolin H even in the absence of TriliE, suggesting that TriliE is not essential for ilicicolin H formation. This observation is in contrast to our results from homologous expression, where the absence of TriliE led to a drastic reduction in ilicicolin H levels. Additionally, *Shenouda et al.*^14^ detected several acetylated shunt products that we – in our homologous system – could neither see in the corresponding extracted ion current chromatograms (EICCs), nor did they appear in the molecular network search. Further dissimilarities were observed in the accumulated intermediates upon TriliE knockout. Whereas *Zhang* et al.^12^ reported an increase in 8-*epi*-ilicicolin and tetramic acid in their system, we observed accumulation of 8-*epi*-ilicicolin and the *bis*-diene intermediate. This variation may be attributed to the more efficient activity of the cytochrome P450 enzyme (TriliC) in our homologous host system, illustrating how foreign hosts can influence product formation in different ways. In accordance with the other studies, we confirmed that TriliA is indispensable to initiate ilicicolin formation, underscoring its critical role in the pathway.

By applying molecular networking to our untargeted metabolomics data, we identified several features related to ilicicolin H and its pathway intermediates. NMR analysis of one of these compounds, which exhibited an unusual modification, revealed a novel ilicicolin product that we have named ilicicolin K. Initial antimycotic tests showed that ilicicolin K possesses antifungal activity. Further studies will be conducted to evaluate its effectiveness against pathogenic fungal strains.

## References

(1) Hayakawa, S.; Minato, H.; Katagiri, K. THE ILICICOLINS, ANTIBIOTICS FROM CYLINDROCLADIUM ILICICOLA. J Antibiot (Tokyo) 1971, 24 (9), 653–654. 10.7164/antibiotics.24.653.

(2) Singh, S. B.; Liu, W.; Li, X.; Chen, T.; Shafiee, A.; Card, D.; Abruzzo, G.; Flattery, A.; Gill, C.; Thompson, J. R.; Rosenbach, M.; Dreikorn, S.; Hornak, V.; Meinz, M.; Kurtz, M.; Kelly, R.; Onishi, J. C. Antifungal Spectrum, in Vivo Efficacy, and Structure-Activity Relationship of Ilicicolin h. ACS Med Chem Lett 2012, 3 (10), 814–817. 10.1021/ML300173E.

(3) Gutierrez-Cirlos, E. B.; Merbitz-Zahradnik, T.; Trumpower, B. L. Inhibition of the Yeast Cytochrome Bc1 Complex by Ilicicolin H, a Novel Inhibitor That Acts at the Qn Site of the Bc1 Complex. Journal of Biological Chemistry 2004, 279 (10), 8708–8714. 10.1074/jbc.M311805200.

(4) Guzmán, E. A.; Pitts, T. P.; Tandberg, K. R.; Winder, P. L.; Wright, A. E.; Guzmán, E. A. ; Pitts, T. P. ; Tandberg, K. R. ; Winder, P. L. ;; Wright, A. E. Discovery of Survivin Inhibitors Part 1: Screening the Harbor Branch Pure Compound Library. Marine Drugs 2021, Vol. 19, Page 73 2021, 19 (2), 73. 10.3390/MD19020073.

(5) Luo, X.; Huang, B.; Lu, H.; Zhang, Y.; Gan, X.; Wang, X.; Liu, Y. Bioactive Alkaloids from the Beibu Gulf Coral-Associated Fungus Acremonium Sclerotigenum GXIMD 02501. Records of Natural Products 2022, 17 (1), 165–169. 10.25135/rnp.329.2204.2437.

(6) Li, M.; Zhang, A.; Qi, X.; Yu, R.; Li, J. A Novel Inhibitor of PGK1 Suppresses the Aerobic Glycolysis and Proliferation of Hepatocellular Carcinoma. Biomedicine & Pharmacotherapy 2023, 158, 114115. 10.1016/j.biopha.2022.114115.

(7) Singh, S. B.; Liu, W.; Li, X.; Chen, T.; Shafiee, A.; Dreikorn, S.; Hornak, V.; Meinz, M.; Onishi, J. C. Structure–Activity Relationship of Cytochrome Bc1 Reductase Inhibitor Broad Spectrum Antifungal Ilicicolin H. Bioorg Med Chem Lett 2013, 23 (10), 3018–3022. 10.1016/J.BMCL.2013.03.023.

(8) Singh, S. B.; Li, X.; Chen, T. Biotransformation of Antifungal Ilicicolin H. Tetrahedron Lett 2011, 52 (46), 6190–6191. 10.1016/J.TETLET.2011.09.051.

(9) Matsumoto, M.; Minato, H. Structure of Ilicocolin H, an Antifungal Antibiotic. Tetrahedron Lett 1976, 17 (42), 3827–3830. 10.1016/S0040-4039(00)93121-6.

(10) Tanabe, M.; Urano, S. Biosynthetic Studies with 13C. Tetrahedron 1983, 39 (21), 3569–3574. 10.1016/S0040-4020(01)88667-1.

(11) Junker, B.; Zhang, J.; Mann, Z.; Reddy, J.; Greasham, R. Scale-Up Studies on a Defined Medium Process for Pilot Plant Production of Illicicolin by Gliocladium Roseum. Biotechnol Prog 2001, 17 (2), 278–286. 10.1021/bp0001718.

(12) Zhang, Z.; Jamieson, C. S.; Zhao, Y. L.; Li, D.; Ohashi, M.; Houk, K. N.; Tang, Y. Enzyme-Catalyzed Inverse-Electron Demand Diels-Alder Reaction in the Biosynthesis of Antifungal Ilicicolin H. J Am Chem Soc 2019, 141 (14), 5659–5663. 10.1021/jacs.9b02204.

(13) Lin, X.; Yuan, S.; Chen, S.; Chen, B.; Xu, H.; Liu, L.; Li, H.; Gao, Z. Heterologous Expression of Ilicicolin H Biosynthetic Gene Cluster and Production of a New Potent Antifungal Reagent, Ilicicolin J. Molecules 2019, 24 (12), 2267. 10.3390/molecules24122267.

(14) Shenouda, M. L.; Ambilika, M.; Cox, R. J. Trichoderma Reesei Contains a Biosynthetic Gene Cluster That Encodes the Antifungal Agent Ilicicolin H. Journal of Fungi 2021, 7 (12), 1034. 10.3390/jof7121034.

(15) Gilchrist, C. L. M.; Chooi, Y. H. Clinker & Clustermap.Js: Automatic Generation of Gene Cluster Comparison Figures. Bioinformatics 2021, 37 (16), 2473–2475. 10.1093/BIOINFORMATICS/BTAB007.

(16) Tomico-Cuenca, I.; Mach, R. L.; Mach-Aigner, A. R.; Derntl, C. An Overview on Current Molecular Tools for Heterologous Gene Expression in Trichoderma. Fungal Biology and Biotechnology 2021 8:1 2021, 8 (1), 1–17. 10.1186/S40694-021-00119-2.

(17) Mandels, M.; Andreotti, R. E. Problems and Challenges in the Cellulose to Cellulase Fermentation. Process Biochem.; (United Kingdom) 1978, 13:5.

(18) Steiger, M. G.; Vitikainen, M.; Uskonen, P.; Brunner, K.; Adam, G.; Pakula, T.; Penttilä, M.; Saloheimo, M.; MacH, R. L.; Mach-Aigner, A. R. Transformation System for Hypocrea Jecorina (Trichoderma Reesei) That Favors Homologous Integration and Employs Reusable Bidirectionally Selectable Markers. Appl Environ Microbiol 2011, 77 (1), 114–121. 10.1128/AEM.02100-10/FORMAT/EPUB.

(19) Derntl, C.; Kiesenhofer, D. P.; Mach, R. L.; Mach-Aigner, A. R. Novel Strategies for Genomic Manipulation of Trichoderma Reesei with the Purpose of Strain Engineering. Appl Environ Microbiol 2015, 81 (18), 6314–6323. 10.1128/AEM.01545-15.

(20) Derntl, C.; Mach, R. L.; Mach-Aigner, A. R. Fusion Transcription Factors for Strong, Constitutive Expression of Cellulases and Xylanases in Trichoderma Reesei. Biotechnol Biofuels 2019, 12 (1), 231. 10.1186/s13068-019-1575-8.

(21) Schiestl, R. H.; Gietz, R. D. High Efficiency Transformation of Intact Yeast Cells Using Single Stranded Nucleic Acids as a Carrier. Curr Genet 1989, 16 (5–6), 339–346. 10.1007/BF00340712/METRICS.

(22) Christianson, T. W.; Sikorski, R. S.; Dante, M.; Shero, J. H.; Hieter, P. Multifunctional Yeast High-Copy-Number Shuttle Vectors. Gene 1992, 110 (1), 119–122. 10.1016/0378-1119(92)90454-W.

(23) Punt, P. J.; Oliver, R. P.; Dingemanse, M. A.; Pouwels, P. H.; van den Hondel, C. A. M. J. J. Transformation of Aspergillus Based on the Hygromycin B Resistance Marker from Escherichia Coli. Gene 1987, 56 (1), 117–124. 10.1016/0378-1119(87)90164-8.

(24) Pfaffl, M. W. A New Mathematical Model for Relative Quantification in Real-Time RT–PCR. Nucleic Acids Res 2001, 29 (9), e45–e45. 10.1093/NAR/29.9.E45.

(25) Steiger, M. G.; Mach, R. L.; Mach-Aigner, A. R. An Accurate Normalization Strategy for RT-QPCR in Hypocrea Jecorina (Trichoderma Reesei). J Biotechnol 2010, 145 (1), 30–37. 10.1016/J.JBIOTEC.2009.10.012.

(26) Skowronek, P.; Thielert, M.; Voytik, E.; Tanzer, M. C.; Hansen, F. M.; Willems, S.; Karayel, O.; Brunner, A. D.; Meier, F.; Mann, M. Rapid and In-Depth Coverage of the (Phospho-)Proteome With Deep Libraries and Optimal Window Design for Dia-PASEF. Molecular & Cellular Proteomics 2022, 21 (9), 100279. 10.1016/J.MCPRO.2022.100279.

(27) Sikorski, R. S.; Hieter, P. A System of Shuttle Vectors and Yeast Host Strains Designed for Efficient Manipulation of DNA in Saccharomyces Cerevisiae. Genetics 1989, 122 (1), 19–27. 10.1093/genetics/122.1.19.

(28) Dührkop, K.; Fleischauer, M.; Ludwig, M.; Aksenov, A. A.; Melnik, A. V.; Meusel, M.; Dorrestein, P. C.; Rousu, J.; Böcker, S. SIRIUS 4: A Rapid Tool for Turning Tandem Mass Spectra into Metabolite Structure Information. Nat Methods 2019, 16 (4), 299–302. 10.1038/s41592-019-0344-8.

(29) Hoffmann, M. A.; Nothias, L.-F.; Ludwig, M.; Fleischauer, M.; Gentry, E. C.; Witting, M.; Dorrestein, P. C.; Dührkop, K.; Böcker, S. Assigning Confidence to Structural Annotations from Mass Spectra with COSMIC. bioRxiv 2021, 2021.03.18.435634. 10.1101/2021.03.18.435634.

(30) Dührkop, K.; Shen, H.; Meusel, M.; Rousu, J.; Böcker, S. Searching Molecular Structure Databases with Tandem Mass Spectra Using CSI:FingerID. Proceedings of the National Academy of Sciences 2015, 112 (41), 12580–12585. 10.1073/pnas.1509788112.

(31) Schmid, R.; Heuckeroth, S.; Korf, A.; Smirnov, A.; Myers, O.; Dyrlund, T. S.; Bushuiev, R.; Murray, K. J.; Hoffmann, N.; Lu, M.; Sarvepalli, A.; Zhang, Z.; Fleischauer, M.; Dührkop, K.; Wesner, M.; Hoogstra, S. J.; Rudt, E.; Mokshyna, O.; Brungs, C.; Ponomarov, K.; Mutabdžija, L.; Damiani, T.; Pudney, C. J.; Earll, M.; Helmer, P. O.; Fallon, T. R.; Schulze, T.; Rivas-Ubach, A.; Bilbao, A.; Richter, H.; Nothias, L. F.; Wang, M.; Orešič, M.; Weng, J. K.; Böcker, S.; Jeibmann, A.; Hayen, H.; Karst, U.; Dorrestein, P. C.; Petras, D.; Du, X.; Pluskal, T. Integrative Analysis of Multimodal Mass Spectrometry Data in MZmine 3. Nature Biotechnology 2023 41:4 2023, 41 (4), 447–449. 10.1038/s41587-023-01690-2.

(32) Nothias, L. F.; Petras, D.; Schmid, R.; Dührkop, K.; Rainer, J.; Sarvepalli, A.; Protsyuk, I.; Ernst, M.; Tsugawa, H.; Fleischauer, M.; Aicheler, F.; Aksenov, A. A.; Alka, O.; Allard, P. M.; Barsch, A.; Cachet, X.; Caraballo-Rodriguez, A. M.; Da Silva, R. R.; Dang, T.; Garg, N.; Gauglitz, J. M.; Gurevich, A.; Isaac, G.; Jarmusch, A. K.; Kameník, Z.; Kang, K. Bin; Kessler, N.; Koester, I.; Korf, A.; Le Gouellec, A.; Ludwig, M.; Martin H, C.; McCall, L. I.; McSayles, J.; Meyer, S. W.; Mohimani, H.; Morsy, M.; Moyne, O.; Neumann, S.; Neuweger, H.; Nguyen, N. H.; Nothias-Esposito, M.; Paolini, J.; Phelan, V. V.; Pluskal, T.; Quinn, R. A.; Rogers, S.; Shrestha, B.; Tripathi, A.; van der Hooft, J. J. J.; Vargas, F.; Weldon, K. C.; Witting, M.; Yang, H.; Zhang, Z.; Zubeil, F.; Kohlbacher, O.; Böcker, S.; Alexandrov, T.; Bandeira, N.; Wang, M.; Dorrestein, P. C. Feature-Based Molecular Networking in the GNPS Analysis Environment. Nature Methods 2020 17:9 2020, 17 (9), 905–908. 10.1038/s41592-020-0933-6.

(33) Schmid, R.; Petras, D.; Nothias, L. F.; Wang, M.; Aron, A. T.; Jagels, A.; Tsugawa, H.; Rainer, J.; Garcia-Aloy, M.; Dührkop, K.; Korf, A.; Pluskal, T.; Kameník, Z.; Jarmusch, A. K.; Caraballo-Rodríguez, A. M.; Weldon, K. C.; Nothias-Esposito, M.; Aksenov, A. A.; Bauermeister, A.; Albarracin Orio, A.; Grundmann, C. O.; Vargas, F.; Koester, I.; Gauglitz, J. M.; Gentry, E. C.; Hövelmann, Y.; Kalinina, S. A.; Pendergraft, M. A.; Panitchpakdi, M.; Tehan, R.; Le Gouellec, A.; Aleti, G.; Mannochio Russo, H.; Arndt, B.; Hübner, F.; Hayen, H.; Zhi, H.; Raffatellu, M.; Prather, K. A.; Aluwihare, L. I.; Böcker, S.; McPhail, K. L.; Humpf, H. U.; Karst, U.; Dorrestein, P. C. Ion Identity Molecular Networking for Mass Spectrometry-Based Metabolomics in the GNPS Environment. Nature Communications 2021 12:1 2021, 12 (1), 1–12. 10.1038/s41467-021-23953-9.

(34) Pino, L. K.; Searle, B. C.; Bollinger, J. G.; Nunn, B.; MacLean, B.; MacCoss, M. J. The Skyline Ecosystem: Informatics for Quantitative Mass Spectrometry Proteomics. Mass Spectrom Rev 2020, 39 (3), 229–244. 10.1002/MAS.21540.

(35) Demichev, V.; Messner, C. B.; Vernardis, S. I.; Lilley, K. S.; Ralser, M. DIA-NN: Neural Networks and Interference Correction Enable Deep Proteome Coverage in High Throughput. Nature Methods 2019 17:1 2019, 17 (1), 41–44. 10.1038/s41592-019-0638-x.

(36) Demichev, V.; Szyrwiel, L.; Yu, F.; Teo, G. C.; Rosenberger, G.; Niewienda, A.; Ludwig, D.; Decker, J.; Kaspar-Schoenefeld, S.; Lilley, K. S.; Mülleder, M.; Nesvizhskii, A. I.; Ralser, M. Dia-PASEF Data Analysis Using FragPipe and DIA-NN for Deep Proteomics of Low Sample Amounts. Nature Communications 2022 13:1 2022, 13 (1), 1–8. 10.1038/s41467-022-31492-0.

(37) Tyanova, S.; Temu, T.; Sinitcyn, P.; Carlson, A.; Hein, M. Y.; Geiger, T.; Mann, M.; Cox, J. The Perseus Computational Platform for Comprehensive Analysis of (Prote)Omics Data. Nature Methods 2016 13:9 2016, 13 (9), 731–740. 10.1038/nmeth.3901.

(38) Martinez, D.; Berka, R. M.; Henrissat, B.; Saloheimo, M.; Arvas, M.; Baker, S. E.; Chapman, J.; Chertkov, O.; Coutinho, P. M.; Cullen, D.; Danchin, E. G. J.; Grigoriev, I. V.; Harris, P.; Jackson, M.; Kubicek, C. P.; Han, C. S.; Ho, I.; Larrondo, L. F.; De Leon, A. L.; Magnuson, J. K.; Merino, S.; Misra, M.; Nelson, B.; Putnam, N.; Robbertse, B.; Salamov, A. A.; Schmoll, M.; Terry, A.; Thayer, N.; Westerholm-Parvinen, A.; Schoch, C. L.; Yao, J.; Barbote, R.; Nelson, M. A.; Detter, C.; Bruce, D.; Kuske, C. R.; Xie, G.; Richardson, P.; Rokhsar, D. S.; Lucas, S. M.; Rubin, E. M.; Dunn-Coleman, N.; Ward, M.; Brettin, T. S. Genome Sequencing and Analysis of the Biomass-Degrading Fungus Trichoderma Reesei (Syn. Hypocrea Jecorina). Nature Biotechnology 2008 26:5 2008, 26 (5), 553–560. 10.1038/nbt1403.

(39) Wang, J.; Chitsaz, F.; Derbyshire, M. K.; Gonzales, N. R.; Gwadz, M.; Lu, S.; Marchler, G. H.; Song, J. S.; Thanki, N.; Yamashita, R. A.; Yang, M.; Zhang, D.; Zheng, C.; Lanczycki, C. J.; Marchler-Bauer, A. The Conserved Domain Database in 2023. Nucleic Acids Res 2023, 51 (D1), D384–D388. 10.1093/NAR/GKAC1096.

(40) Marchler-Bauer, A.; Bo, Y.; Han, L.; He, J.; Lanczycki, C. J.; Lu, S.; Chitsaz, F.; Derbyshire, M. K.; Geer, R. C.; Gonzales, N. R.; Gwadz, M.; Hurwitz, D. I.; Lu, F.; Marchler, G. H.; Song, J. S.; Thanki, N.; Wang, Z.; Yamashita, R. A.; Zhang, D.; Zheng, C.; Geer, L. Y.; Bryant, S. H. CDD/SPARCLE: Functional Classification of Proteins via Subfamily Domain Architectures. Nucleic Acids Res 2017, 45 (D1), D200–D203. 10.1093/NAR/GKW1129.

(41) Koike, H.; Aerts, A.; Labutti, K.; Grigoriev, I. V.; Baker, S. E. Comparative Genomics Analysis of Trichoderma Reesei Strains. https://home.liebertpub.com/ind 2013, 9 (6), 352–367. 10.1089/IND.2013.0015.

(42) Lah, L.; Kraševec, N.; Trontelj, P.; Komel, R. High Diversity and Complex Evolution of Fungal Cytochrome P450 Reductase: Cytochrome P450 Systems. Fungal Genetics and Biology 2008, 45 (4), 446–458. 10.1016/j.fgb.2007.10.004.

