## Supplementary Material for "Discovery of a novel antifungal compound, ilicicolin K, through genetic activation of the ilicicolin biosynthetic pathway in *Trichoderma reesei*"

### Table of Contents

#### Supplementary Tables


**Table S1.** Utilized primer for cloning, genotyping and RT-qPCR. S3

**Table S2.** Determined concentrations of ilicicolin H in the culture supernatants of OETriliR and ΔTriliE samples. S4

**Table S3.** Normalization of determined ilicicolin H amounts in the culture supernatant to the respective amount of fungal mycelium in the flasks. S4

**Table S4.** Spectroscopic data of **(3)**: shifts of isolated *bis*-diene. S5

**Table S5.** Spectroscopic data of **(2)**: shifts of isolated 8-*epi*-ilicicolin H. S6

**Table S6.** Spectroscopic data of **(1)**: shifts of isolated ilicicolin H. S7

**Table S7.** Spectroscopic data: shifts of isolated and elucidated ilicicolin K. S9


#### Supplementary Figures


**Figure S1.** Construction of the strain OETriliR. S10

**Figure S2.** Construction of the strain ΔTriliA. S11

**Figure S3.** Construction of the strain ΔTriliE. S12

**Figure S4.** Transcript level analysis using an RT-qPCR assay. S13

**Figure S5.** Determination of the limit of detection (LOD) of ilicicolin H using a matrix matched external calibration curve. S14

**Figure S6.** Matrix matched external calibration curve to quantify ilicicolin H in the medium. S15

**Figure S7.** Histograms of proteomics data distribution per each sample. S16

**Figure S8.** $^{1}$H spectrum (600 MHz) of *bis*-diene **(3)** in acetonitrile-*d*3:DMSO (9:1). S17

**Figure S9.** $^{13}$C spectrum (151 MHz) of *bis*-diene **(3)** in acetonitrile-*d*3:DMSO (9:1). S17

**Figure S10.** $^{1}$H spectrum (600 MHz) of 8-*epi*-ilicicolin H **(2)** in acetonitrile-*d*3. S18

**Figure S11.** $^{13}$C spectrum (151 MHz) of 8-*epi*-ilicicolin H **(2)** in acetonitrile-*d*3. S18

**Figure S12.** $^{1}$H spectrum (600 MHz) of ilicicolin H **(1)** in acetonitrile-*d*3. S19

**Figure S13.** $^{13}$C spectrum (151 MHz) of ilicicolin H **(1)** in acetonitrile-*d*3. S19

**Figure S14.** $^{1}$H spectrum (600 MHz) of ilicicolin K in DMSO-*d*6. S20

**Figure S15.** $^{13}$C spectrum (151 MHz) of ilicicolin K in DMSO-*d*6. S20


#### References: S21

### Supplementary Tables

**Table S1.** Utilized primer for cloning, genotyping and RT-qPCR.

| Name | Sequence | Used for |
|---|---|---|
| 72993_fwd-AflII | CTTAAGCGCAAGAGCATCCACAAATGG | cloning |
| 72993_MRev_SOE | CAACAAATCTCTCTATCAGCGGGCGCAAGTGCTCGTCTCTC | cloning |
| 72993_MFwd_SOE | GAGAGACGAGCACTTGCGCCCGCTGATAGAGAGATTTGTTG | cloning |
| 72993_rev-SpeI | ACTAGTCTAAACAACCGTGCACCGC | cloning |
| 5_ TriliA _fwd-pRS426 | AGTGAGCGCGCGTAATACGACTCACTATAGGGCGAATTGGCAGCACTGTTGGCGATATTC | cloning |
| 5_ TriliA _rev-hph | AGTCACCGGTCACTGTACAGAGCTCGCCTCTAACAATATTCTCTGCTGTG | cloning |
| hph_fwd-5_ TriliA | CACAGCAGAGAATATTGTTAGAGGCGAGCTCTGTACAGTGACCGGTGACT | cloning |
| hph_rev-3_ TriliA | GATATCTGCCGTGCTCTCTAATCGCTCGAGTGGAGATGTGGAGTGGGCGC | cloning |
| 3_ TriliA _fwd-hph | GCGCCCACTCCACATCTCCACTCGAGCGATTAGAGAGCACGGCAGATATC | cloning |
| 3_TriliA_rev-pRS426 | AGAACTAGTGGATCCCCCGGGCTGCAGGAATTCGATATCACTAAGCGCCACTCTAACAGAAG | cloning |
| PgpdA_fwd | GAGCTCTGTACAGTGACCGG | Cloning |
| hph_MR | CGTCAGGACATTGTTGGAGC | cloning / genotyping |
| hph_MF | GACCTGCCTGAAACCGAAC | cloning / genotyping |
| TtrpC_rev | AGGTCGAGTGGAGATGTGGAG | cloning |
| TriliE_5fwd | CAGCCATCATCGCAAACCTAG | cloning |
| TriliE _5rev-hph | CTCCACATCTCCACTCGACCTGTAATGACCATGCCCCAACC | cloning |
| TriliE _3fwd-hph | CCGGTCACTGTACAGAGCTCAGGCGTGGTATCTAGGGCAC | cloning |
| TriliE _3rev | GAGAAGAGAAACGGAGGACTG | cloning |
| 5pyr4_fwd3 | CCAGACGGTGATTCACATATACG | genotyping |
| Ptef_rev-BspTI | CTTAAGTGTGATGTAGCGTGAGAGCTG | genotyping |
| pyr4_3fwd3 | TGCCTTTATCCACATGACGC | genotyping |
| Tpyr4_rev2 | CAGGAAGCTCAGCGTCGAG | genotyping |
| 58285_fwd | GTATGCGTCCTGTGTCTCAC | genotyping |
| 58285_rev | CTGTTACGCAAAGGTCTCCG | genotyping |
| iliE_5'_genomic | AGGTAGTGAAGTCTCGGTTG | genotyping |
| iliE_3'_genomic | CTGAGCTCTGGTATGTGGTT | genotyping |
| sar1fw | TGGATCGTCAACTGGTTCTACGA | RT-qPCR |
| sar1rev | GCATGTGTAGCAACGTGGTCTTT | RT-qPCR |
| act1f | TGAGAGCGGTGGTATCCACG | RT-qPCR |
| act1r | GGTACCACCAGACATGACAATGTTG | RT-qPCR |
| 58285_q1f | GCTGAACATGATGACCAAAGAG | RT-qPCR |

| | | |
|---|---|---|
| 58285_q1r | TTCGTTTTCAGGGAGTAGATGC | RT-qPCR |
| 58289_q1f | AGCAGAAGCCTGGTTTGACTAC | RT-qPCR |
| 58289_q1r | GCTTTGCCGAACTGTGTAGG | RT-qPCR |
| 58953_q1f | CGTCAGAGATGAGGAGGAACTG | RT-qPCR |
| 58953_q1r | GGAATCGTTGAGGCAAGCAC | RT-qPCR |
| Rut75073_q1f | TTGTCAAAGGAGATGATGTCG | RT-qPCR |
| Rut75073_q1r | TACTACAACGTCGACGAAACC | RT-qPCR |
| 76204_q1f | CCGGAAATTCTCGTTGATAGG | RT-qPCR |
| 76204_q1r | ATTGAGCAGGCTCCATTTGTC | RT-qPCR |
| 72993_q1f | CGTTAACGGCTCTTCGGATAC | RT-qPCR |
| 72993_q1r | GCAGAAAAGGAACTCGCGTG | RT-qPCR |

**Table S2.** Determined concentrations of ilicicolin H in the culture supernatants of OETriliR and ΔTriliE samples. *: significant outlier according to Grubbs' test, significance level: 0.05 (two-sided), Z-value: 1.48919.

| | c [ng µL$^{-1}$] | $\overline{X}$ (c, n = 4) [ng µL$^{-1}$] | ± s (c, n = 4) [ng µL$^{-1}$] |
|---|---|---|---|
| **OETriliR #1** | 13.872 | | |
| **OETriliR #2** | 55.553* | 25.295, | 20.319 |
| **OETriliR #3** | 13.128 | (15.208 excl. outlier) | (2.982 excl. outlier) |
| **OETriliR #4** | 18.625 | | |
| **ΔTriliE #1** | 4.251 | | |
| **ΔTriliE #2** | 3.915 | 3.125 | 1.230 |
| **ΔTriliE #3** | 2.806 | | |
| **ΔTriliE #4** | 1.529 | | |

**Table S3.** Normalization of determined ilicicolin H amounts in the culture supernatant to the respective amounts of fungal mycelium in the flasks. *: significant outlier according to Grubbs' test, significance level: 0.05 (two-sided), Z-value: 1.49909.

| | Mycelium amount [mg] | Determined concentration of iliH in the medium [ng µL$^{-1}$] | Absolute amount of ilicicolin H in medium [ng] | [ng] iliH in medium per [mg] mycelium |
|---|---|---|---|---|
| **OETriliR #1** | 208.4 | 13.872 | 1387.220 | 6.657 |
| **OETriliR #2** | 234.3 | 55.553 | 5555.313 | 23.710* |
| **OETriliR #3** | 192.3 | 13.128 | 1312.778 | 6.827 |
| **OETriliR #4** | 304.1 | 18.625 | 1862.527 | 6.125 |
| **ΔTriliE #1** | 187.2 | 4.251 | 425.102 | 2.271 |
| **ΔTriliE #2** | 241.2 | 3.915 | 391.524 | 1.623 |
| **ΔTriliE #3** | 219.6 | 2.806 | 280.648 | 1.278 |
| **ΔTriliE #4** | 180.2 | 1.529 | 152.898 | 0.848 |

**Table S4.** Spectroscopic data of **(3)**: shifts of isolated *bis*-diene.

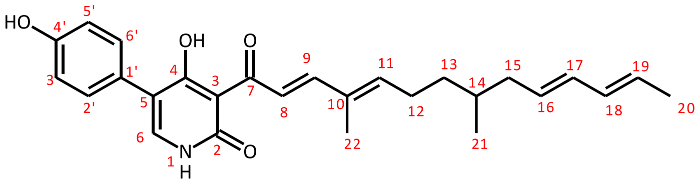

bis-diene **(3)**

| POS | Reported (in CDCl3:Acetone 3:1)[1] | | Found (in ACN:DMSO) | |
|---|---|---|---|---|
| | $\delta_C$ | $\delta_H$(MULT.,INT, *J*) | $\delta_C$ | $\delta_H$(MULT.,INT, *J*) |
| 2 | 162.9 | - | 162.9 | - |
| 3 | 106.7 | - | 106.9 | - |
| 4 | 178.0 | - | 178.2 | - |
| 5 | 114.8 | - | 114.1[c] | - |
| 6 | 139.1 | 7.44 (s, 1H) | 140.6 | 7.41 (s, 1H) |
| 7 | 194.6 | | 194.8 | |
| 8 | 123.3 | 8.01 (d, 1H, *J* = 15.5 Hz) | 123.7 | 8.03 (d, 1H, *J* = 15.46) |
| 9 | 150.1 | 7.53 (d, 1H, *J* = 15.5 Hz) | 159.8 | 7.53 (d, 1H, *J* = 15.4 Hz) |
| 10 | 134.5 | | 134.5 | |
| 11 | 144.8 | 6.01 (t, 1H, *J* = 7.5 Hz) | 145.5 | 6.11 (t, 1H, *J* = 7.27 Hz) |
| 12 | 26.8 | 2.22 (m, 2H) | 26.9 | 2.27 (m, 2H) |
| 13 | 35.5 | Ha 1.22 (m, 1H)<br>Hb 1.42 (m, 1H) | 35.8 | Ha 1.27[b]<br>Hb 1.43 (m, 1H) |
| 14 | 33.1 | 1.50 (m, 1H) | 33.2 | 1.53 (m, 1H) |
| 15 | 40.0 | Ha 1.88 (q, 1H, *J* = 13.0, 7.0 Hz)<br>Hb 2.02 (q, 1H, *J* = 13.0, 6.0 Hz) | 39.9[d] | Ha 1.90[a]<br>Hb 2.05[b] |
| 16 | 130.0 | 5.45 (m, 1H) | 130.7 | 5.55 (m, 1H) |
| 17 | 131.4 | 5.88 (m, 1H) | 132.17 or 132.20 | 6.00 (m, 1H) |
| 18 | 131.6 | 5.94 (m, 1H) | 132.17 or 132.20 | 6.00 (m, 1H) |
| 19 | 126.7 | 5.50 (m, 1H) | 127.3 | 5.55 (m, 1H) |
| 20 | 17.9 | 1.65 (d, 3H, *J* = 6.5 Hz) | 17.7 | 1.69 (d, 3H, *J* = 6.9 Hz) |
| 21 | 19.4 | 0.85 (d, 3H, *J* = 7.0 Hz) | 19.3 | 0.88 (d, 3H, *J* = 6.7 Hz) |
| 22 | 12.4 | 1.84 (s, 3H) | 12.2 | 1.85 (d, 3H, *J* = 1.1) |
| 1’ | 124.4 | | 124.6 | |
| 2’, 6’ | 130.3 | 7.26 (d, 1H, *J* = 8.5 Hz) | 130.8 | 7.25 (d, 1H, *J* = 8.6 Hz) |
| 3’, 5’ | 115.4 | 6.83 (d, 1H, *J* = 8.5 Hz) | 115.6 | 6.81 (d, 1H, *J* = 8.6 Hz) |
| 4’ | 156.9 | | 157.7 | |

[a] Overlayed by ACN solvent signal
[b] Overlayed by Fatty-acid Signal
[c] Hardly visible in 13C-NMR – HMBC Correlation proves position and nature
[d] Overlayed by DMSO Signal – Position from HSQC Correlation

**Table S5.** Spectroscopic data of **(2)**: shifts of isolated 8-*epi*-ilicicolin H.

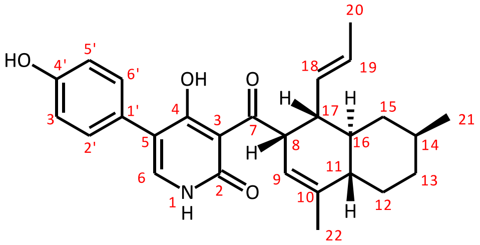


epi-Ilicicolin H **(2)**

| POS | Reported (in acetonitrile-*d*3)[1] $\delta_C$ | $\delta_H$(MULT.,INT, *J*) | Found (in acetonitrile-*d*3) $\delta_C$ | $\delta_H$(MULT.,INT, *J*) |
|---|---|---|---|---|
| 2 | 162.9 | - | 162.45 | - |
| 3 | 108.0 | - | 108.0 | - |
| 4 | 177.6 | - | 177.8 | - |
| 5 | 114.8 | - | 114.5 | - |
| 6 | 140.8 | 7.42 (s, 1H) | 140.8 | 7.40 (s, 1H) |
| 7 | 209.8 | - | 209.9 | - |
| 8 | 50.8 | 5.07 (m, 1H) | 50.7 | 5.07 (m, 1H)[a] |
| 9 | 120.1 | 5.47(m, 1H) | 120.1 | 5.45 (m, 1H) |
| 10 | 141.2 | - | 141.2 | - |
| 11 | 46.0 | 1.54 (m, 1H) | 46.0 | 1.56 (m, 1H)[b] |
| 12 | 30.4 | Ha 1.00 (m, 1H)<br>Hb 2.03 (m, 1H) | 30.5 | Ha 0.98 (m, 1H)<br>Hb 2.06 (m, 1H) |
| 13 | 36.3 | Ha 0.93 (m, 1H)<br>Hb 1.77 (m, 1H) | 36.4 | Ha 0.98 (m, 1H)<br>Hb 1.77 (m, 1H) |
| 14 | 33.7 | 1.41 (m, 1H) | 33.7 | 1.42 (m, 1H) |
| 15 | 40.7 | Ha, 0.44 (q, 1H *J* =24.5 12.0 Hz)<br>Hb 1.74 (m, 1H) | 40.7 | Ha, 0.46 (app. q, 1H *J* =11.8 Hz)<br>Hb 1.77 (m, 1H) |
| 16 | 40.6 | 1.93 (m, 1H) | 40.66 | 1.94 (m, 1H)[c] |
| 17 | 47.4 | 2.17 (m, 1H) | 47.4 | 2.18 (m, 1H) |
| 18 | 133.4 | 5.45 (m, 1H) | 133.5 | 5.45 (m, 1H) |
| 19 | 127.3 | 5.44 (m, 1H) | 127.3 | 5.45 (m, 1H) |
| 20 | 18.1 | 1.54 (d, 3H) | 18.1 | 1.56 (d, 3H, *J* = 4.58 Hz) |
| 21 | 23.0 | 0.88 (d, 3H, *J* = 6.5 Hz) | 23.03 | 0.90 (d, 3H, *J* = 6.6 Hz) |
| 22 | 21.5 | 1.66 (s, 3H) | 21.5 | 1.68 (d, 3H, *J* = 1.6 Hz) |
| 1’ | 125.4 | - | 125.5 | - |
| 2’, 6’ | 131.4 | 7.26 (d, 1H, *J* = 9.0 Hz) | 131.5 | 7.26 (d, 1H, *J* = 8.6 Hz) |
| 3’, 5’ | 116.0 | 6.83 (d, 1H, *J* = 9.0 Hz) | 116.0 | 6.83 (d, 1H, *J* = 8.6 Hz) |
| 4’ | 157.5 | - | 157.5 | - |

[a] Peakshape identical to Literature, small coupling between H8 – H17 indicates cis configuration.
[b] Peak overlayed by H20 – shift from HSQC
[c] Peak overlayed by residual solvent signal – shift from HSQC

**Table S6.** Spectroscopic data of **(1)**: shifts of isolated ilicicolin H.

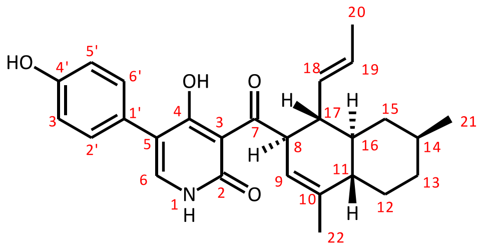


Ilicicolin H **(1)**

| POS | Reported (in acetonitrile-*d*3)[1] | | Found (in acetonitrile-*d*3) | |
|---|---|---|---|---|
| | δC | δH(MULT.,INT, *J*) | δC | δH(MULT.,INT, *J*) |
| 2 | 162.8 | - | 162.9 | - |
| 3 | 107.8 | - | 107.8 | - |
| 4 | 177.6 | - | 177.5 | - |
| 5 | 114.6 | - | 114.3 | - |
| 6 | 140.9 | 7.43 (s,1H) | 141.2 | 7.41 (s, 1H) |
| 7 | 210.7 | - | 210.7 | - |
| 8 | 53.8 | 4.98 (m, 1H) | 53.7 | 4.98 (m, 1H)[a] |
| 9 | 120.4 | 5.21 (m, 1H) | 120.6 | 5.21 (m, 1H) |
| 10 | 139.1 | - | 139.0 | - |
| 11 | 45.1 | 1.67 (m, 1H) | 45.2 | 1.69 (1H, m) |
| 12 | 30.6 | Ha 0.97 (m, 1H) | 30.6 | Ha 0.98 (1H, m) |
| | | Hb 2.03 (m, 1H) | | Hb 2.05 (1H, m) |
| 13 | 36.2 | Ha 0.97 (m, 1H) | 36.2 | Ha 0.98 (1H, m) |
| | | Hb 1.76 (m, 1H) | | Hb 1.78 (1H, m) |
| 14 | 33.4 | 1.38 (m, 1H) | 33.4 | 1.39 (m, 1H) |
| 15 | 40.7 | Ha 0.57 (q, 1H) | 40.4 | Ha 0.58 (dt, 1H, *J* = 13.1, 11.7) |
| | | Hb 1.76 (m, 1H) | | Hb 1.78 (1H, m) |
| 16 | 44.2 | 1.22 (m, 1H) | 44.2 | 1.23 (m, 1H) |
| 17 | 45.8 | 2.48 (m, 1H) | 45.8 | 2.49 (td, 1H, *J* = 10.8, 9.1 Hz) |
| 18 | 134.2 | 5.21 (m, 1H) | 134.4 | 5.22 (m, 1H) |
| 19 | 126.8 | 5.32 (m, 1H) | 126.8 | 5.33 (dq, 1H, *J* = 15.3, 6.3 Hz) |
| 20 | 18.0 | 1.52 (dd, 3H, *J* = 6.5, 1.3 Hz ) | 18.1 | 1.53 (dd, 3H, *J* = 6.3, 1.6 Hz) |
| 21 | 23.0 | 0.88 (d, 3H, *J* = 6.5 Hz) | 23.0 | 0.89 (d, 3H, *J* = 6.6 Hz) |
| 22 | 21.0 | 1.62 (s, 3H) | 21.1 | 1.64 (1H, m) |
| 1’ | 125.3 | - | 125.0 | - |
| 2’, 6’ | 131.4 | 7.26 (d, 1H, *J* = 8.6 Hz) | 131.3 | 7.27 (d, 1H, *J* = 8.6 Hz) |
| 3’, 5’ | 116.0 | 6.83 (d, 1H, *J* = 8.6 Hz) | 116.1 | 6.84 (d, 1H, *J* = 8.6 Hz) |
| 4’ | 157.5 | - | 158.1 | - |

[a] Peakshape in accordance with literature

Structure elucidation of ilicicolin K was mostly based on established assignments from ilicicolin H **(1)**, and relative changes of the recorded spectra found. Noticeable the aliphatic region of the molecule stayed unaltered according to NMR. Peak shapes as well as shifts remained comparable to **(1)**. The biggest differences were found in the aromatic region, where the former para-substituted phenyl ring exhibited two uncoupled signals with intensity of 1H each. We therefore assumed that the phenyl ring had to be substituted at four positions. One of the two new substitutions can be explained by the addition of a hydroxy group at position C3' . $^{1}$H-NMR shows three acidic protons, two of which show HMBC correlations to aromatic carbon signals that can be attributed to two phenolic OHs groups on the aromatic ring. Especially the general shift of all aromatic carbons in $^{13}$C-NMR towards highfield is an indication for electron donating effects into the phenyl ring. The other substitution was hypothesized to be a link of C6' to the OH at C4 of **(1)**. A direct proof of this new link via NMR was impossible due to the lack of protons in the relevant region. Advanced NMR techniques to clarify the carbon skeleton (like INADEQUATE or ADEQUATE) were not feasible with the amount of material at hand. However, the resonance structure of C4 in **(1)** gives C4 a partially carbonylic nature, which is seen in the very far downfield shift of that signal (177 ppm). Etherification of the hydroxyl group at C4 would lead to a stabilization the enol, thus a significantly lower chemical shift. In the structure of ilicicolin K one finds a shift of 16 ppm upfield for that specific signal, to 161 ppm. These values are in the range where *para*-hydroxypyridines and their respective ethers are usually found[2]. Therefore, direct link between the aromatic ring and the hydroxy-group of the former C4 of **(1)** is very likely.

The resulting structure fits to the found mass of 447 m/z and is shown below. Assignment of this structure based on $^{1}$H, $^{13}$C, COSY, HSQC and HMBC spectra was possible and in compliance with expected signal positions for such a structure.

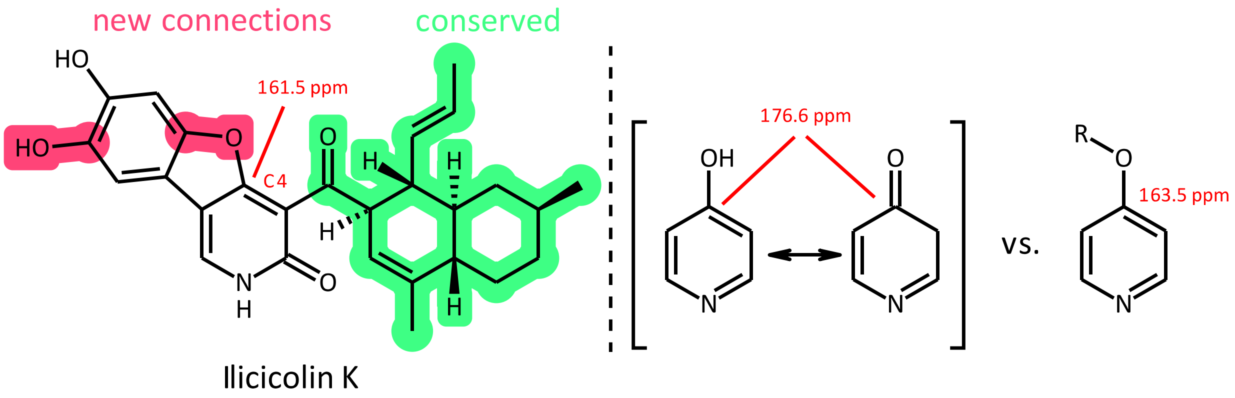


**Left:** Elucidated structure of the newly found Ilicicolin family member. **Right:** $^{13}$C-NMR shift of 4-pyrodinone and the alkylated derivate found in literature[2].

**Table S7.** Spectroscopic data: shifts of isolated and elucidated ilicicolin K. The numbering for positions in Ilicicolin K was changed compared to Ilicicolin H **(1)**, as the parent system now is fluorene. Since, compared to **(1)**, all carbon positions can be directly mapped, the numbers that would correspond to **(1)** are also indicated.

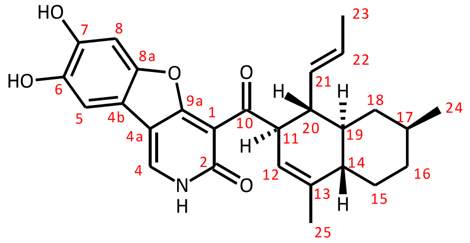


Ilicicolin K

| Position in Ilicicolin K | Position in **(1)** | Found (in DMSO-*d6*)[1] δC | δH(MULT.,INT, *J*) | Comparison to **(1)** δC | delta |
|---|---|---|---|---|---|
| 1 | 3 | 110.2 | -- | 107.8 | 2.4 |
| 2 | 2 | 165.0 | -- | 162.9 | 2.1 |
| 4 | 6 | 131.6 | 8.30 (s, 1H) | 141.2 | -9.6 |
| 4a | 5 | 110.6 | -- | 114.3 | -3.7 |
| 4b | 1' | 111.7 | -- | 125.0 | -13.3 |
| 5 | 2' | 99.1 | 6.91 (s, 1H) | 131.3 | -32.2 |
| 6 | 3' | 143.4[a] | -- | 116.1 | 27.3 |
| 7 | 4' | 146.4[a] | -- | 158.1 | -11.7 |
| 8 | 5' | 106.6 | 7.24 (s, 1H) | 116.1 | -9.5 |
| 8a | 6' | 150.8 | -- | 131.3 | 19.5 |
| 9a | 4 | 161.5 | -- | 177.5 | -16 |
| 10 | 7 | 201.0 | -- | 210.7 | -9.7 |
| 11 | 8 | 54.2 | 4.36 (ddt, 1H, *J* = 10.3, 4.1, 2.4 Hz) | 53.7 | 0.5 |
| 12 | 9 | 120.6 | 5.24 (m, 1H) | 120.6 | 0 |
| 13 | 10 | 137.2 | -- | 139.0 | -1.83 |
| 14 | 11 | 44.2 | 1.60 (m, 1H) | 45.2 | -1 |
| 15 | 12 | 29.8 | Ha 1.99 (m, 1H)<br>Hb 0.92 (m, 1H) | 30.6 | -0.8 |
| 16 | 13 | 35.5 | Ha 1.71 (m, 1H)<br>Hb 0.92 (m, 1H) | 36.2 | -0.7 |
| 17 | 14 | 32.6 | 1.34 (m, 1H) | 33.4 | -0.8 |
| 18 | 15 | 40.5 | Ha 1.71 (m,1H)<br>Hb 051 (app. q, 1H, *J* = 12.1 Hz) | 40.4 | 0.1 |
| 19 | 16 | 43.5 | 1.13 (qd, 1H, *J* = 11.5, 2.8 Hz) | 44.2 | -0.7 |
| 20 | 17 | 46.0 | 2.25 (m, 1H) | 45.8 | 0.2 |
| 21 | 18 | 133.4 | 5.08 (ddd, 1H, J = 15.2, 9.23, 1.7 Hz) | 134.4 | -1 |
| 22 | 19 | 125.9 | 5.24 (m, 1H) | 126.8 | -0.9 |
| 23 | 20 | 18.0 | 1.28 (dd, 3H, J = 6.4, 1.6 Hz) | 18.1 | -0.1 |
| 24 | 21 | 23.1 | 0.85 (d, 3H, J = 6.6 Hz) | 23.0 | 0.1 |
| 25 | 22 | 21.2 | 1.60 (m, 3H) | 21.1 | 0.1 |
| [a] Assignment of C6 and C7 is ambiguous between each other | | | | | |

### Supplementary Figures

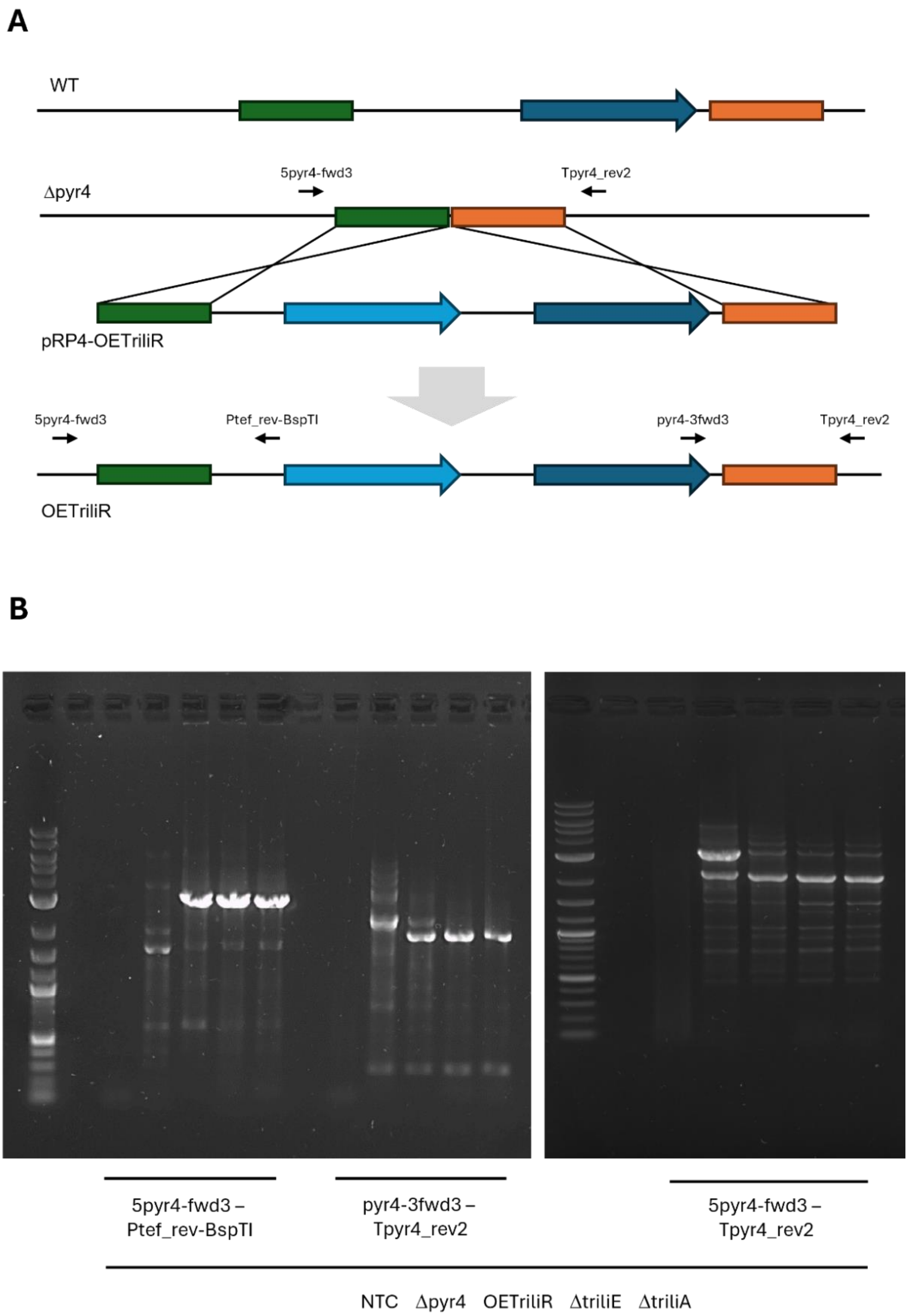


**Figure S1.** Construction of the strain OETriliR. (A) The coding region for the transcription factor TriliR was put under the control of the *tef1* promoter and inserted into the *pyr4* locus of the recipient strain Δpyr4 using the strategy described by *Derntl et al.*[3]. (B) The insertion of the TriliR expression cassette was tested on a genomic level by suitable PCR assays using the indicated primers and the chromosomal DNA of the used strains. NTC, no template control.

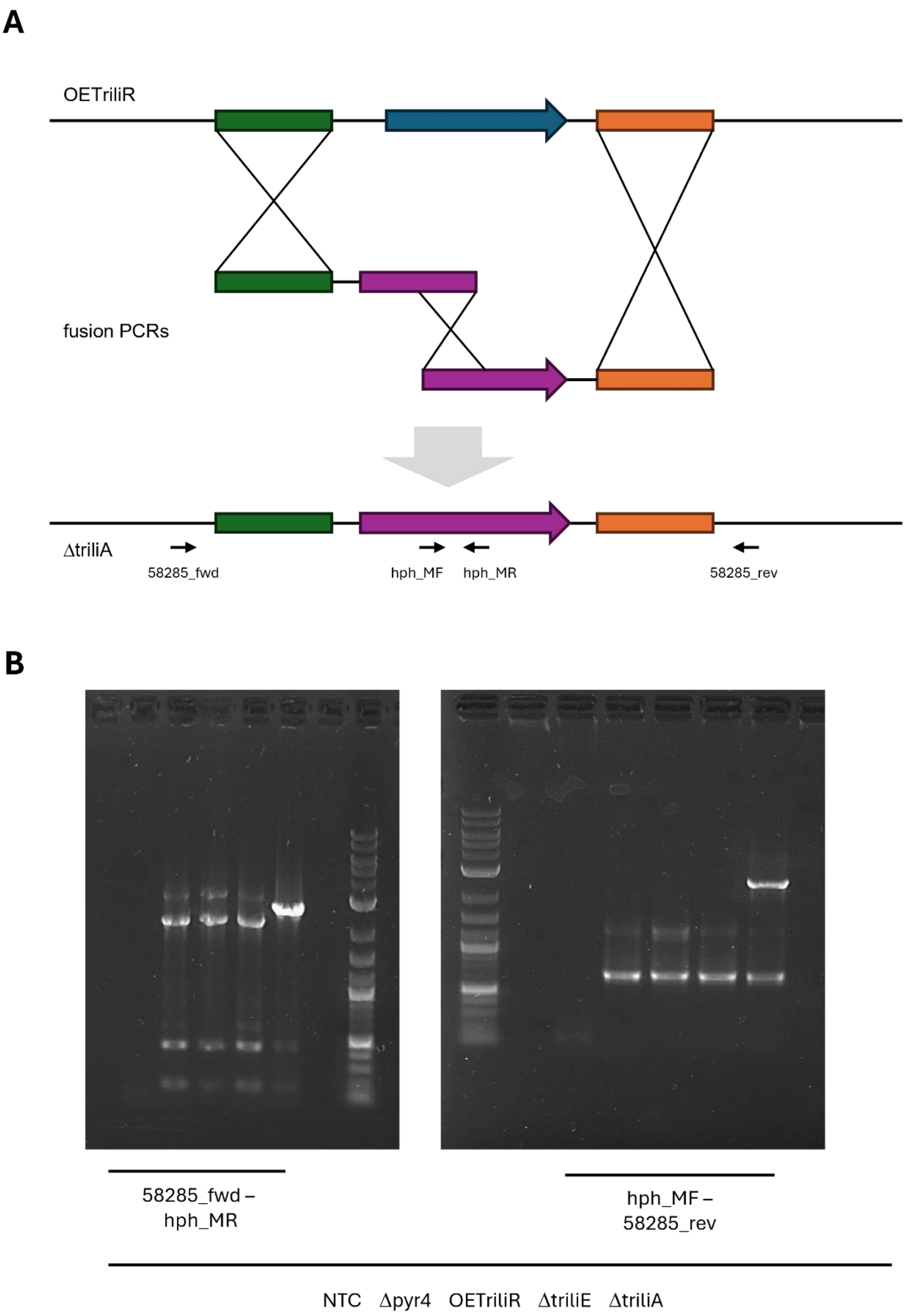


**Figure S2.** Construction of the strain ΔTriliA. (A) To delete the gene *triliA*, a split marker strategy, and the hygromycin resistance marker *hph* were used. (B) The deletion was tested on a genomic level by suitable PCR assays using the indicated primers and the chromosomal DNA of the used strains. NTC, no template control.

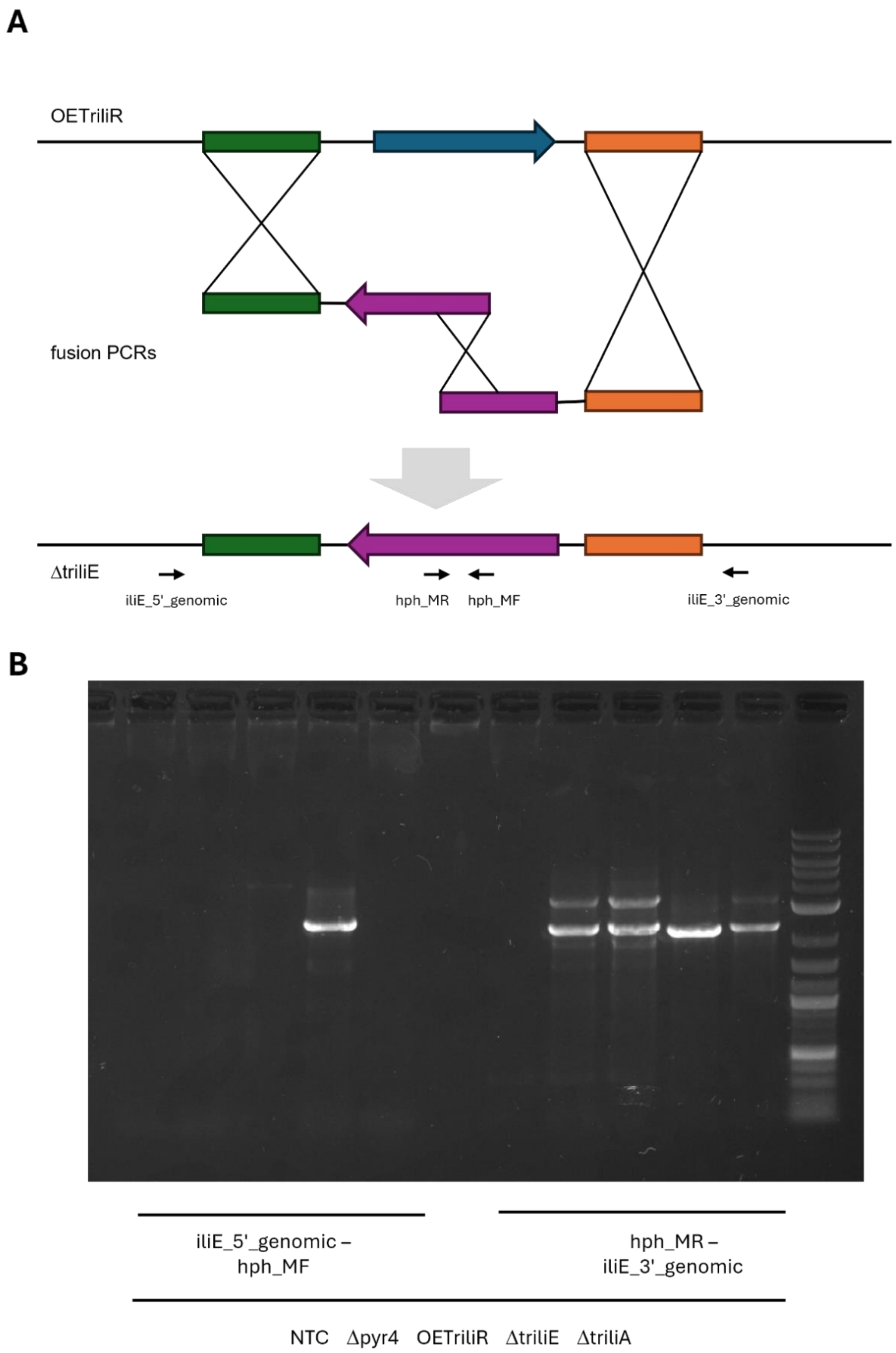


**Figure S3.** Construction of the strain ΔTriliE. (A) To delete the gene *triliE*, a split marker strategy, and the hygromycin resistance marker *hph* were used. (B) The deletion was tested on a genomic level by suitable PCR assays using the indicated primers and the chromosomal DNA of the used strains. NTC, no template control.

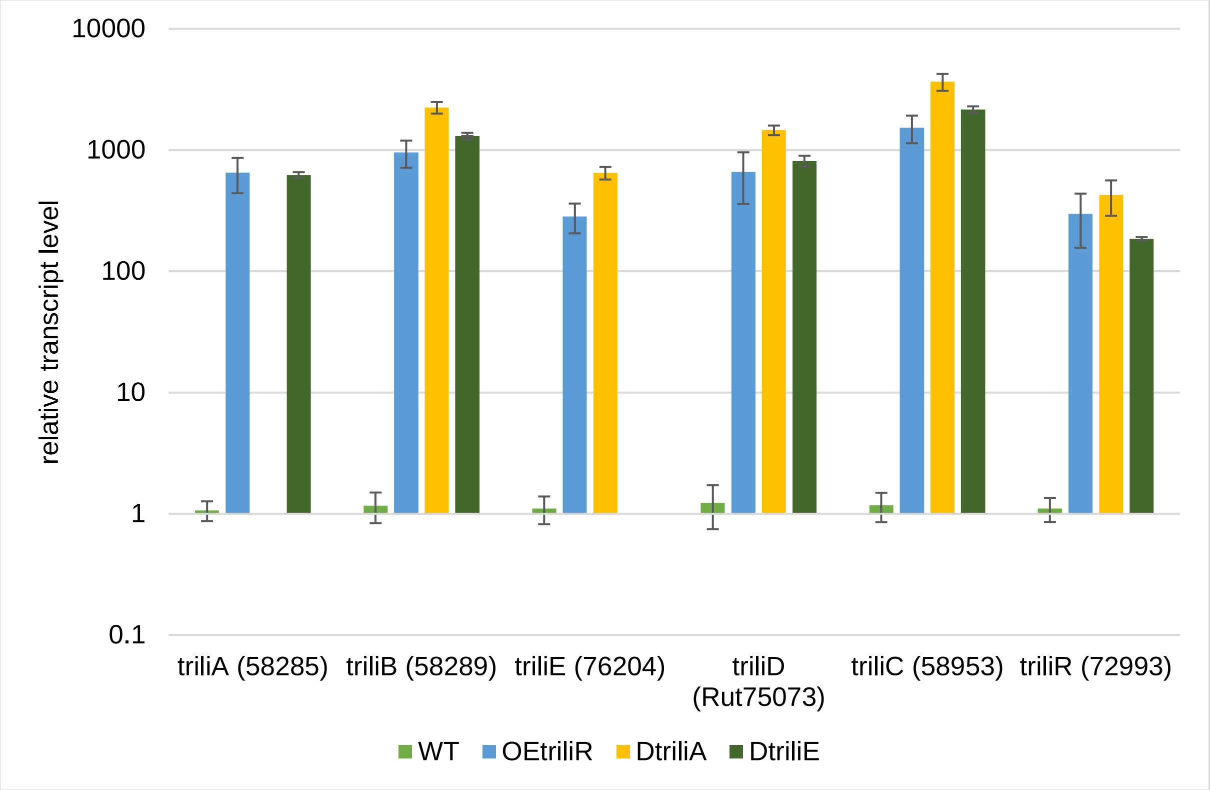


**Figure S4.** Transcript level analysis using an RT-qPCR assay. The *T. reesei* strains QM6a Δmus53 (wildtype, WT), OETriliR, ΔTriliA, and ΔTriliE were cultivated in MAM with glycerol as carbon source in biological quadruplicates. After 72 hours, the total RNA was extracted, and the cDNA was synthesized using the NEB LunaScript kit. The cDNAs were used as template for the RT-qPCR reactions using the NEB LunaMix according to the manufacturer's instructions. The relative transcript levels of the indicated genes were normalized to the average value of the wildtype samples (WT, QM6a Δmus53) using the Pfaffl[4] method and the genes *sar1* and *act1* as reference genes. Error bars indicate standard deviation.

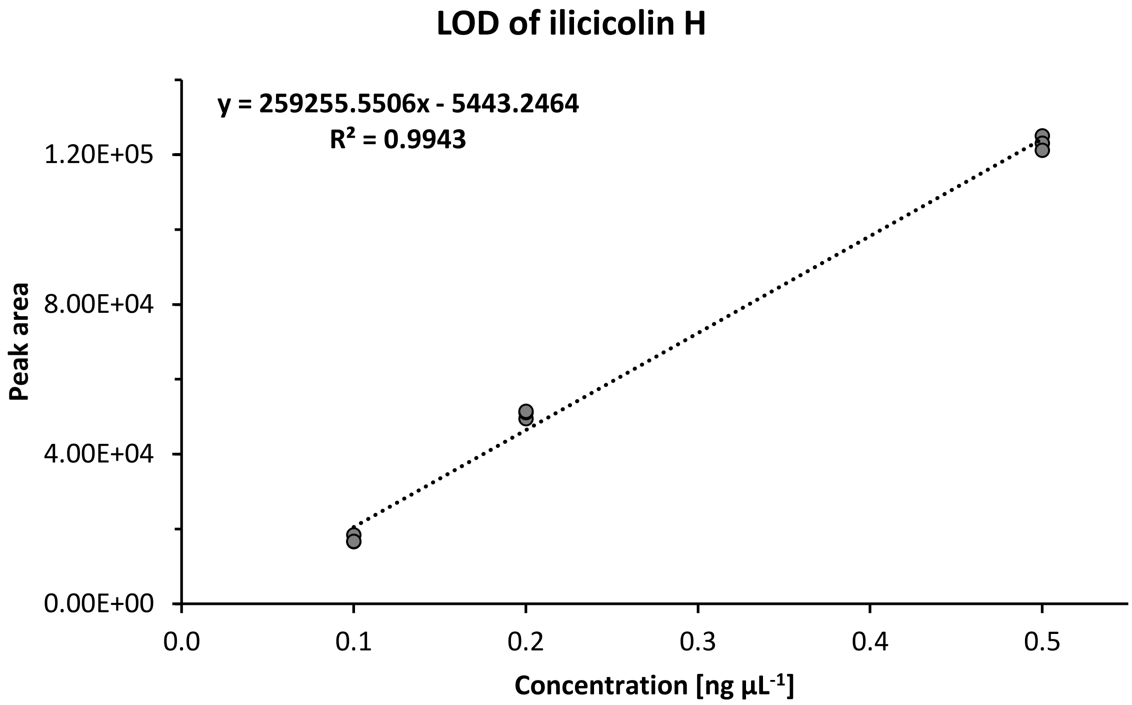


**Figure S5.** Determination of the limit of detection (LOD) of ilicicolin H using a matrix matched external calibration curve. The timsTOF Pro Mass Spectrometer was set to detect signals with an intensity threshold of I = 100, classifying signals below this threshold as noise. In the extracted ion current chromatograms (EICCs) of ilicicolin H, no signals (defined as having an intensity < 100) were detected in the wildtype and ΔTriliA mycelial extracts. Given that the mycelia contain the compound at higher concentrations, it is reasonable to assume that the media of the wildtype and ΔTriliA strains are practically free of ilicicolin H, due to the dilution effect of the large volume of the medium. Consequently, the media from the wildtype and ΔTriliA were pooled, and low amounts of ilicicolin H standard were spiked into it, to create matrix-matched standard solutions with a concentration of 0.1, 0.2 and 0.5 ng µL$^{-1}$. The standards were measured in technical triplicates. The resulting peak areas were plotted against the known concentrations to generate a linear regression curve. The derived equation (y = 259255.5506x - 5443.2464) was set equal to the intensity threshold of I = 100, yielding a limit of detection of 0.02138 ng µL$^{-1}$ (= 21.38 ng mL$^{-1}$).

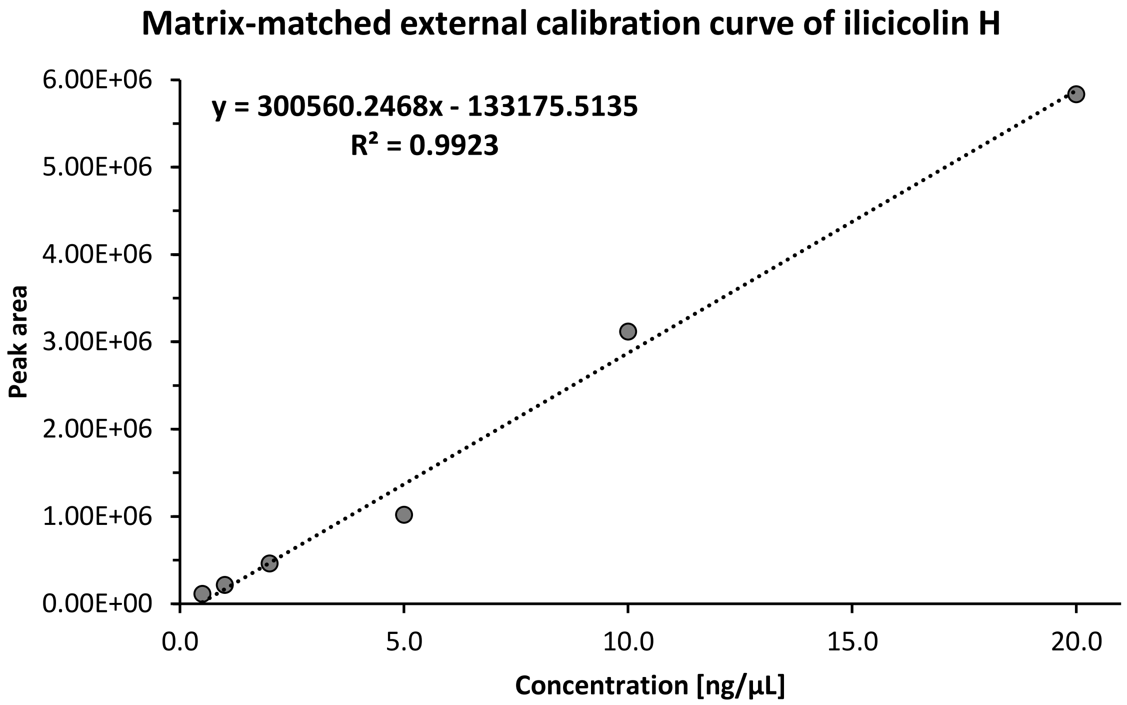


**Figure S6.** Matrix matched external calibration curve to quantify ilicicolin H in the medium. As matrix, wildtype and ΔTriliA media were pooled. Using the obtained equation (y = 300560.2468x - 133175.5135), the concentration of ilicicolin H was calculated in the samples. (see **Table S2** and **Table S3**).

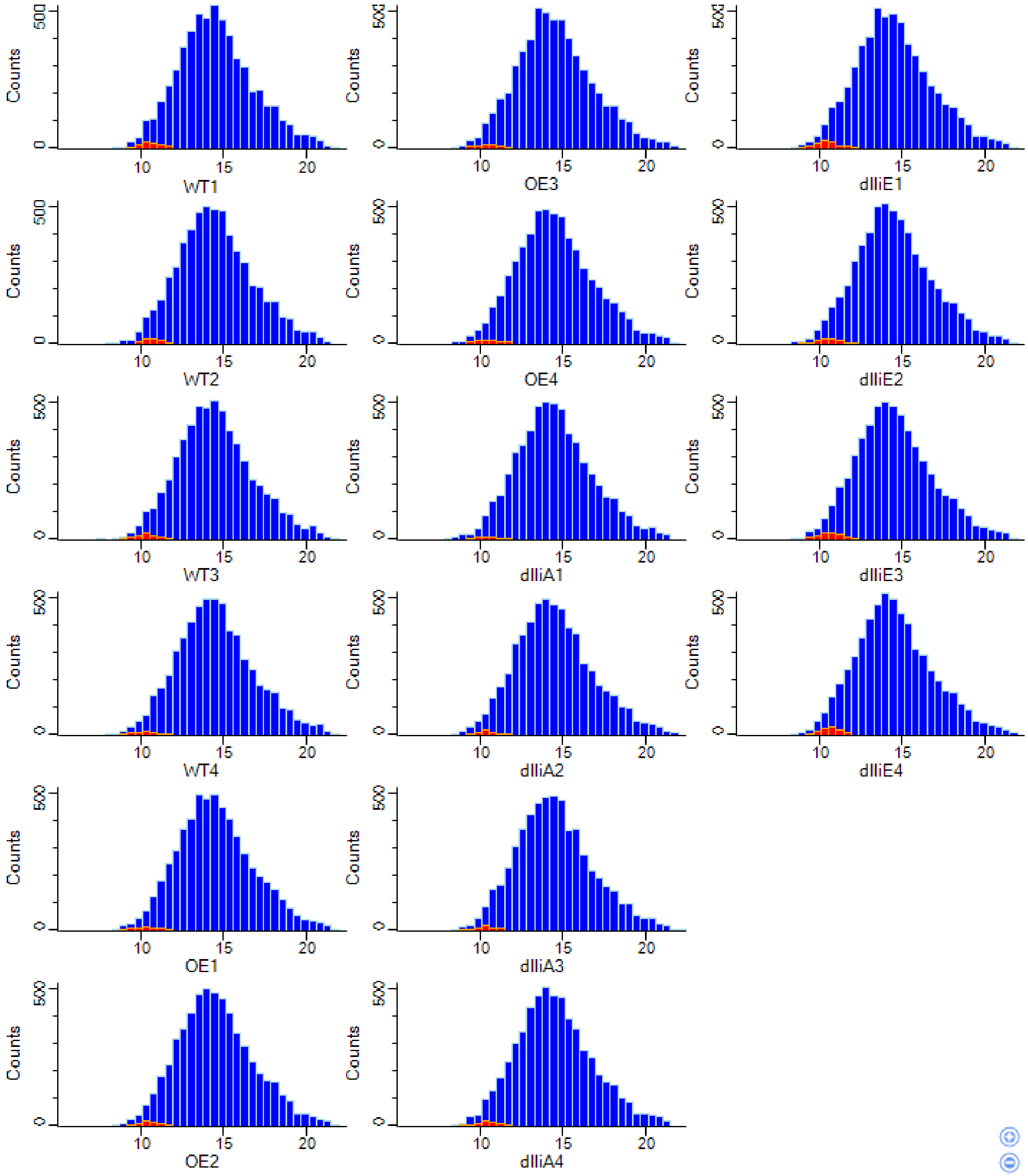


**Figure S7.** Histograms of proteomics data distribution per each sample. Imputed values are highlighted in red. Proteins were filtered for 4 valid values in at least one group. Imputation of missing values was conducted based on normal distribution (width 0.3, down shift 1.8).

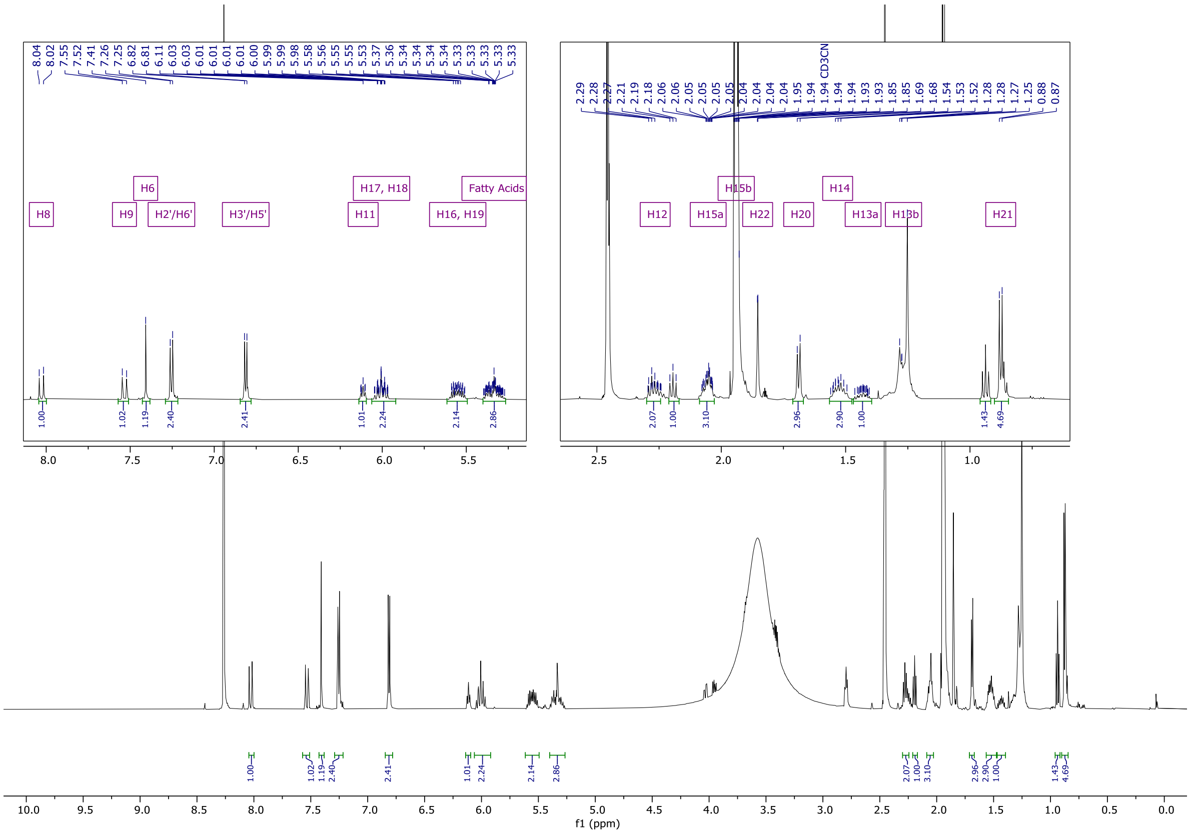


**Figure S8**. $^{1}$H spectrum (600 MHz) of *bis*-diene **(3)** in acetonitrile-*d*3:DMSO (9:1).

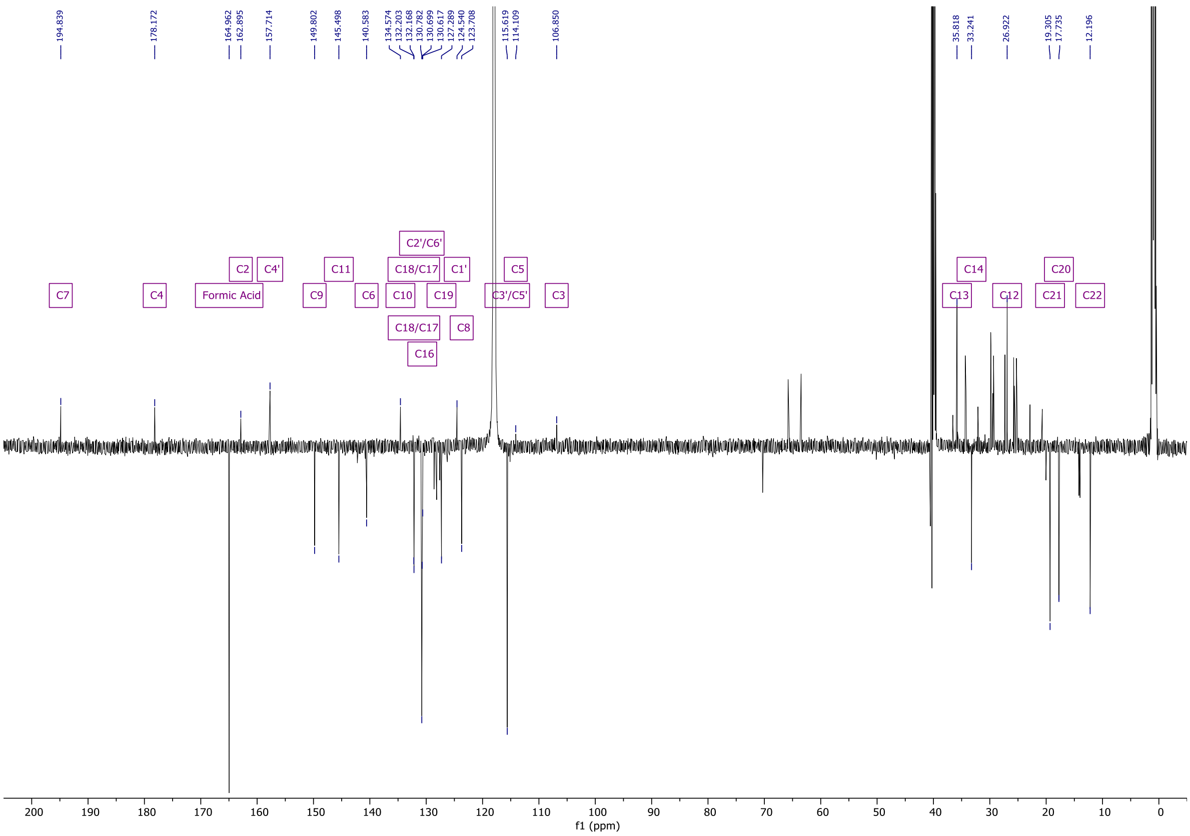


**Figure S9.** $^{13}$C spectrum (151 MHz) of *bis*-diene **(3)** in acetonitrile-*d*3:DMSO (9:1).

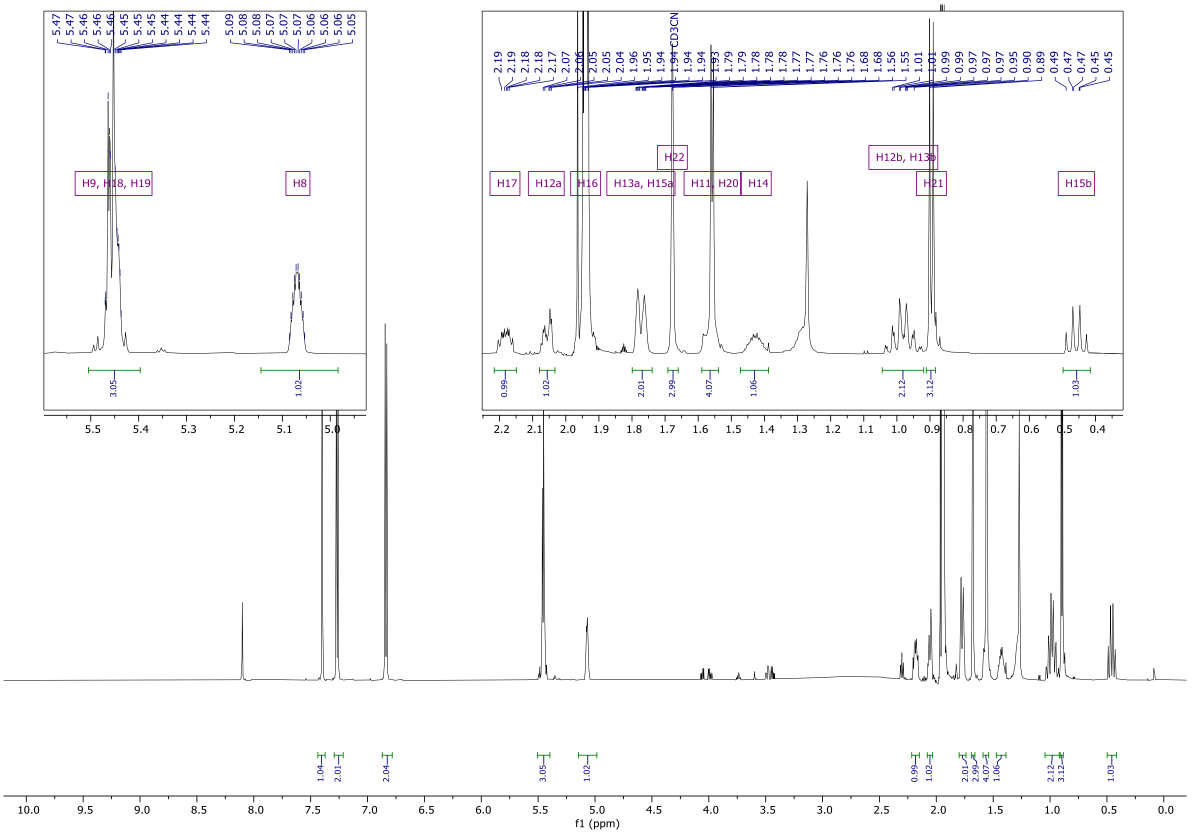


**Figure S10.** $^{1}H$ spectrum (600 MHz) of 8-*epi*-ilicicolin H **(2)** in acetonitrile-*d*3.

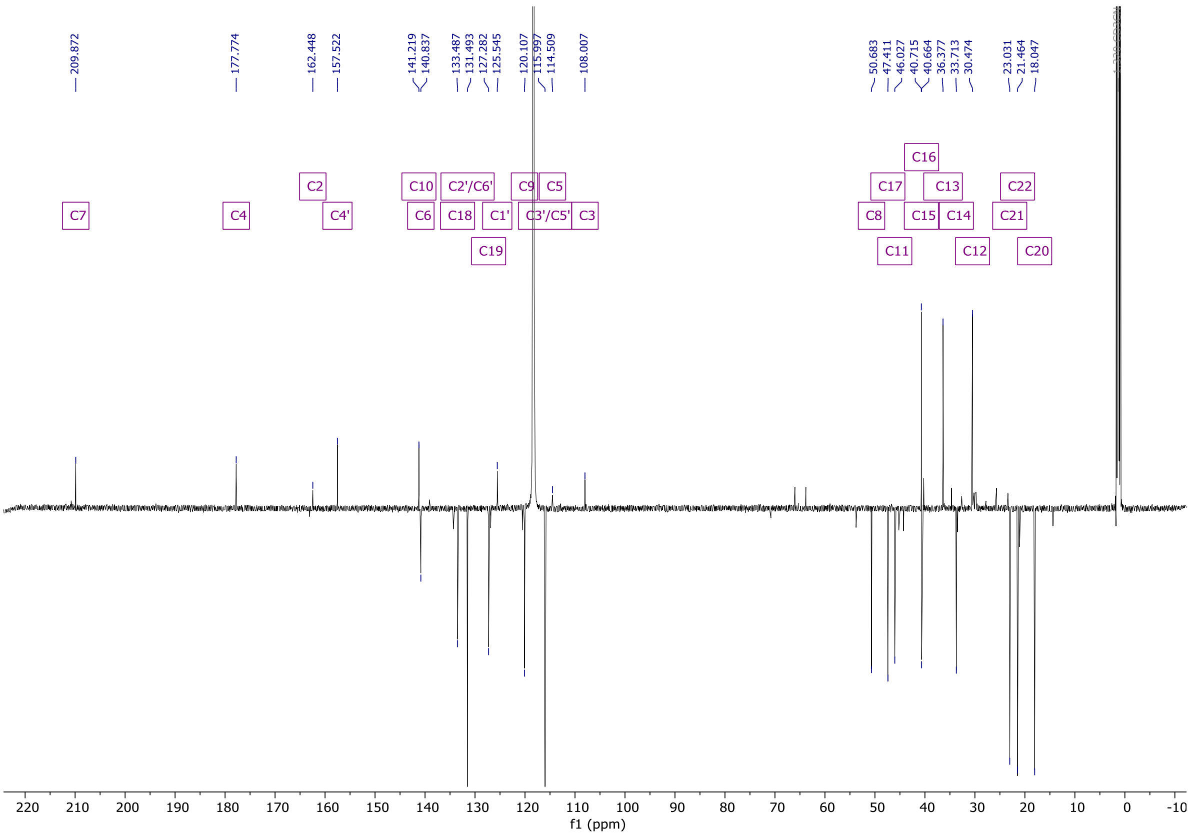


**Figure S11.** $^{13}C$ spectrum (151 MHz) of 8-*epi*-ilicicolin H **(2)** in acetonitrile-*d*3.

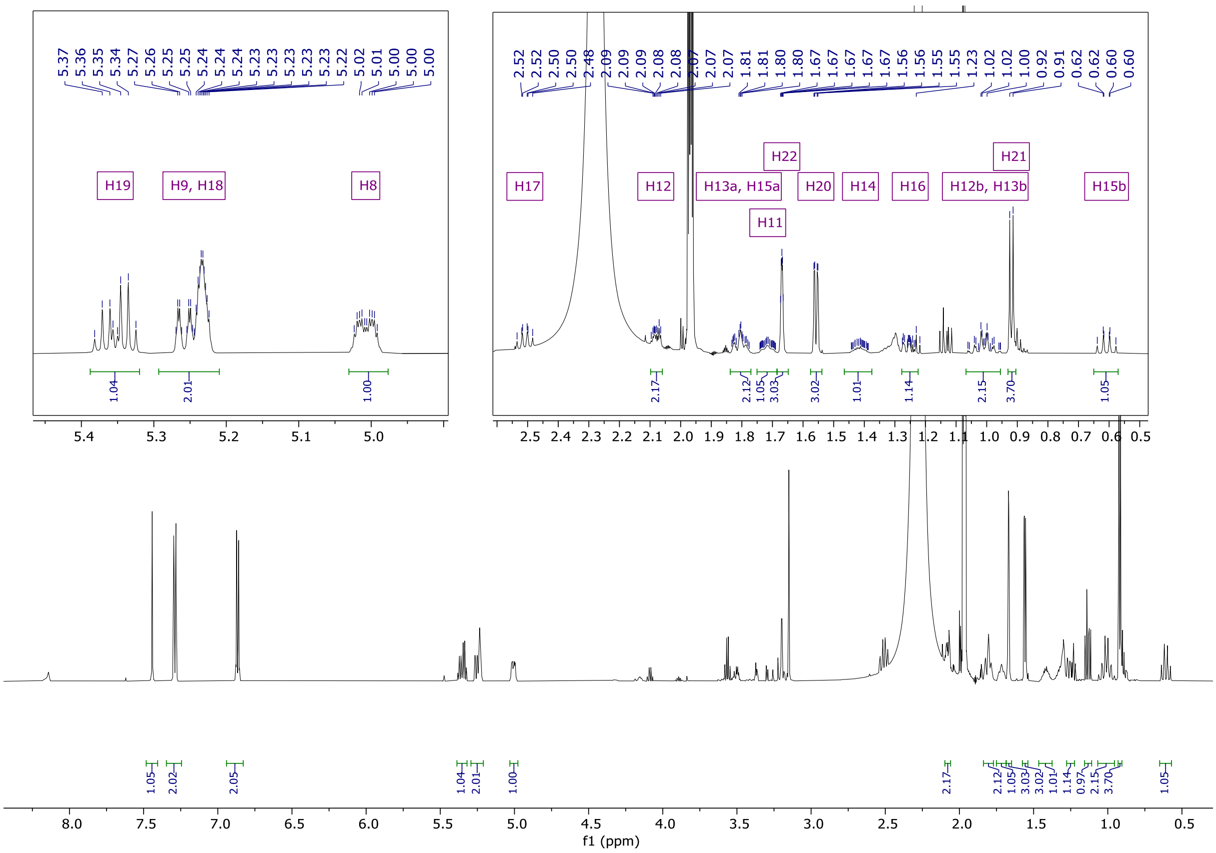


**Figure S12.** $^{1}$H spectrum (600 MHz) of ilicicolin H **(1)** in acetonitrile-*d*3.

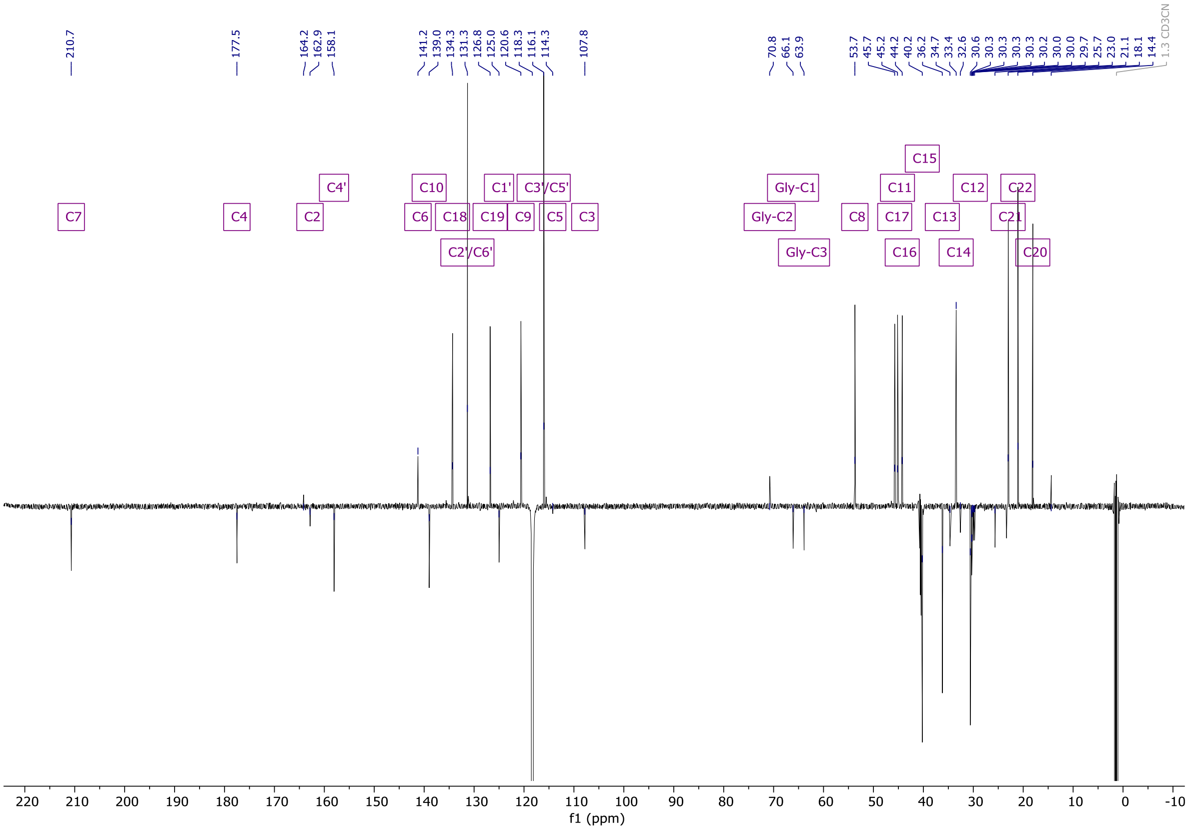


**Figure S13.** $^{13}$C spectrum (151 MHz) of ilicicolin H **(1)** in acetonitrile-*d*3.

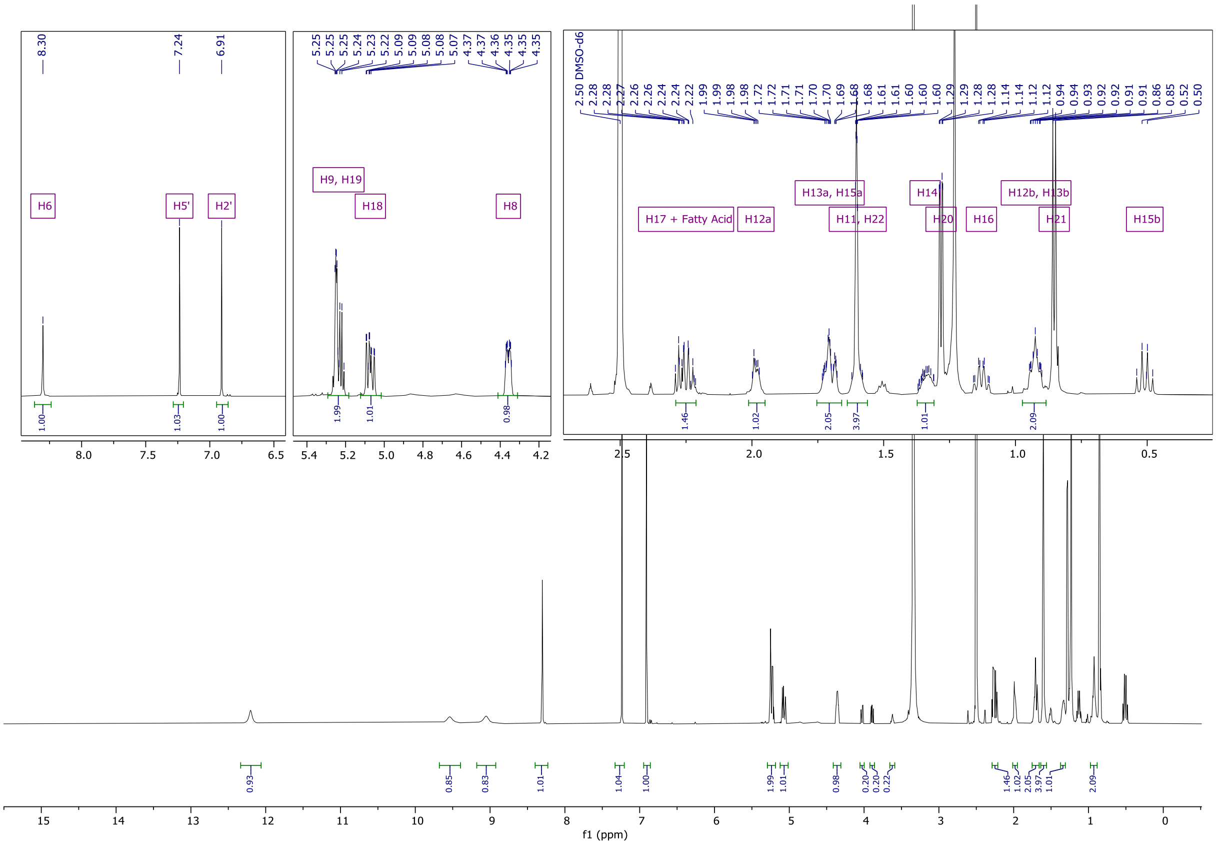


**Figure S14.** $^{1}$H spectrum (600 MHz) of ilicicolin K in DMSO-*d*6.

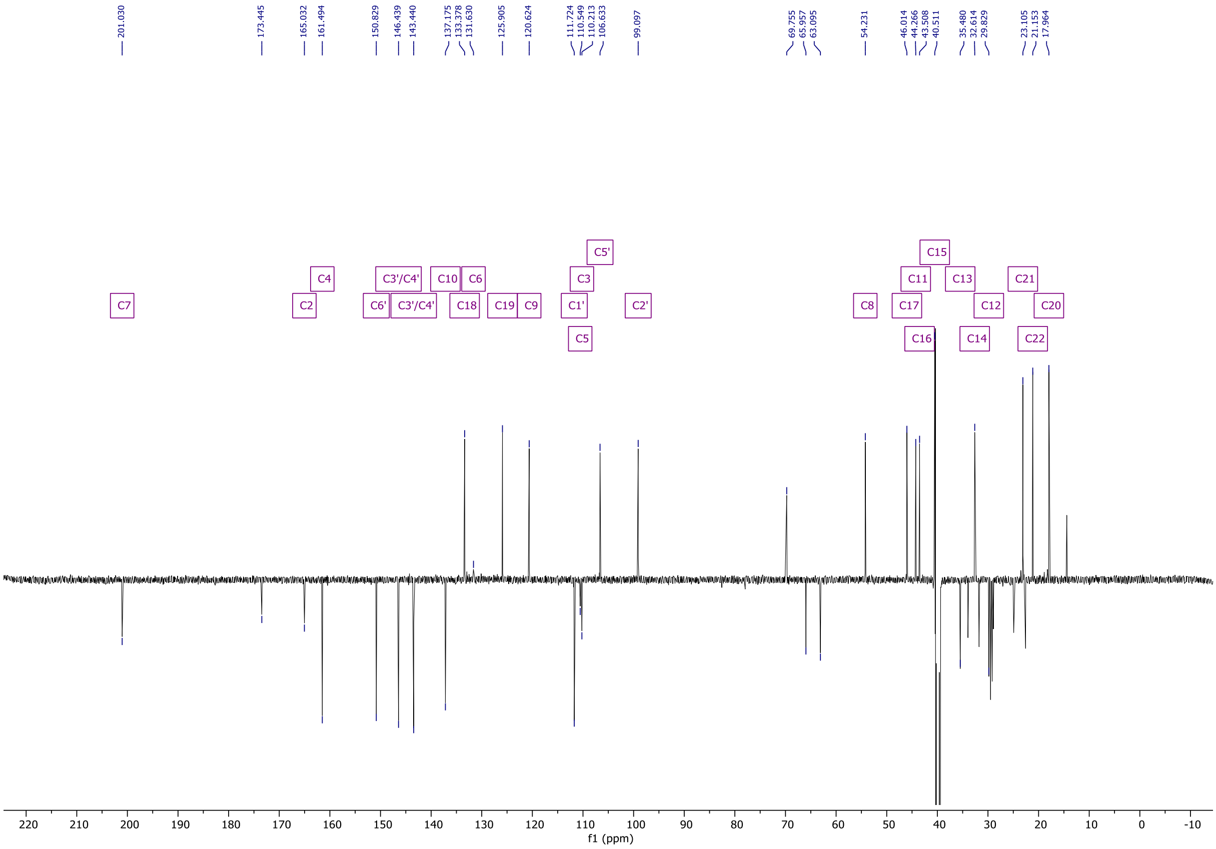


**Figure S15.** $^{13}$C spectrum (151 MHz) of ilicicolin K in DMSO-*d*6.

### References


(1) Zhang, Z.; Jamieson, C. S.; Zhao, Y. L.; Li, D.; Ohashi, M.; Houk, K. N.; Tang, Y. Enzyme-Catalyzed Inverse-Electron Demand Diels-Alder Reaction in the Biosynthesis of Antifungal Ilicicolin H. *J Am Chem Soc* **2019**, *141* (14), 5659–5663. https://doi.org/10.1021/jacs.9b02204.

(2) Todorov, A. R.; Wirtanen, T.; Helaja, J. Photoreductive Removal of O -Benzyl Groups from Oxyarene N -Heterocycles Assisted by O -Pyridine–Pyridone Tautomerism. *J Org Chem* **2017**, *82* (24), 13756–13767. https://doi.org/10.1021/acs.joc.7b02775.

(3) Derntl, C.; Kiesenhofer, D. P.; Mach, R. L.; Mach-Aigner, A. R. Novel Strategies for Genomic Manipulation of Trichoderma Reesei with the Purpose of Strain Engineering. *Appl Environ Microbiol* **2015**, *81* (18), 6314–6323. https://doi.org/10.1128/AEM.01545-15.

(4) Pfaffl, M. W. A New Mathematical Model for Relative Quantification in Real-Time RT–PCR. *Nucleic Acids Res* **2001**, *29* (9), e45–e45. https://doi.org/10.1093/NAR/29.9.E45.
